# An NLR Integrated Domain toolkit to identify plant pathogen effector targets

**DOI:** 10.1101/2021.08.23.457316

**Authors:** David Landry, Isabelle Mila, Cyrus Raja Rubenstein Sabbagh, Matilda Zaffuto, Cécile Pouzet, Dominique Tremousaygue, Patrick Dabos, Laurent Deslandes, Nemo Peeters

**Author notes:** Author contributions: DLA: designed experiments, performed most experiments, analyzed data, wrote manuscript IMI: performed split-luciferase experiment, analyzed data CSA: cloned the T3E inY2H plasmids MZA: participated in the Y2H screening process CPO: performed the FRET-FLIM experiment, analyzed data TRE: performed Arabidosis-Rs inoculations, analyzed the data PDA: performed Arabidosis-Rs inoculations, analyzed the data LDE: designed experiments, analyzed data, wrote manuscript NPE: designed experiments, analyzed data, wrote manuscript.

## Abstract

Plant resistance genes (or NLR “Nod-like Receptors”) are known to contain atypical domains procuring them with a decoy capacity. Some of these integrated domains (or ID) allow the plant to lure the virulence determinants (“effectors”) of pathogens and triggering a specific NLR immune reaction.

In this work, our goal was to generate a library of known IDs that could be screened with plant pathogen effectors in order to identify putative new effector virulence targets and NLR-effector pairs.

We curated the IDs contained in NLRs from seven model and crop plant species. We cloned 52 IDs representing 31 distinct Pfam domains. This library was screened for interaction by yeast-two-hybrid with a set of 31 conserved *Ralstonia solanacearum* type III effectors. This screening and the further *in planta* interaction assay allowed us to identify three interactions, involving different IDs (kinase, DUF3542, WRKY) and two type III effectors (RipAE and PopP2).

PopP2 was found to physically interact with ID#85, an atypical WRKY domain integrated in the GmNLR-ID85 NLR protein from Soybean. Using a imaging method in living plant cells, we showed that PopP2 associates with ID#85 in the nucleus. But unlike the known WRKY-containing Arabidopsis RRS1-R NLR receptor, this newly identified soybean WRKY domain could not be acetylated by PopP2 and its atypical sequence (WRKYGKR) also probably renders it inefficient in plant immunity triggering.

This ID toolkit is available for screening with other plant pathogen effectors and should prove useful to discover new effectors targets and potentially engineer new plant resistance genes.

## INTRODUCTION

Plants can resist pathogen attacks thanks to their immune system. To generate a suitable and dedicated defence response, plants must be able to detect microbes and discriminate between friend or foe [1]. The plant immunity relies on two main types of immune receptors: extracellular pattern recognition receptors (PRRs) and intracellular Nod-like receptor (NLR) [2, 3]. PRRs monitor the extracellular environment and detect distinct evolutionarily conserved structures, known as pathogen-associated molecular patterns (PAMPs). Upon PAMPs recognition, PRRs trigger activation of basal immunity also called PAMP-triggered immunity (PTI). This basal defence is generally sufficient to prevent the plants from being colonized by pathogens [4]. Throughout evolution, successful pathogens, able to cause disease on a host, have evolved sophisticated means to defeat PTI. In most cases, these are secreted or translocated proteins acting as virulence factors (effectors). In plant pathogenic bacteria, the type III secretion system (T3SS) enables the injection of type III effectors (T3Es) inside the host cell in order to subvert basal defence responses or to manipulate the host metabolism [5, 6]. Resistant plants are genetically equipped to perceive specific T3Es. Here, NLRs recognize directly or indirectly the presence or the activities of matching T3Es [3, 7]. Such perception activates a strong immune response known as effector-triggered immunity (ETI) that is often associated with a localized cell death called the hypersensitive response (HR) [2, 3].

The plant NLR family belongs to the STAND (signal-transduction ATPases with numerous domains) P-loop ATPases of the AAA+ superfamily [8]. These immune receptors have a tripartite modular architecture [1]. NLRs possess conserved domains including a C-terminal leucine-rich repeats (LRR) domain, a central nucleotide-binding (NB) domain and a variable N-terminal domain. Usually, NLRs contains a Toll/Interleukin-1 receptor (TIR) or Coiled-Coil (CC) domain at their N-terminus extremity defining two major classes of NLR proteins, the TNLs and CNLs, respectively [9]. Recently, the presence of atypical integrated domains (“ID” hereafter) in NLRs (NLR-ID) have also been reported [10-12]. Comparative analyses of plant NLR architecture suggests that the integration of these atypical domains is a widespread phenomenon occurring in all plant lineages, but only in a limited number of NLRs. In addition, some of these unconventional NLR domains can be homologous to host targets of pathogen effectors suggesting a potential role as decoy [11]. IDs can occur in NLR singleton but are also found in paired NLR [10-12]. NLR pairs are encoded by two genes in head-to-head orientation with a common promoter region suggesting their coregulation [13]. In several cases, NLR pair forms an oligomeric complex in which each member has a distinct role: one detects the pathogen (“sensor”) while the other induces the defence responses (“executor”). The latter sees its signaling activity repressed by the sensor and derepressed in the presence of specific virulence factor [14, 15]. The ID function has mainly been documented in paired NLRs, representing a new recognition model known as the “integrated decoy” model [16, 17]. The ID in the sensor NLR acts as a decoy that lures the effector and diverts it from its real virulence target. Upon targeting of the integrated decoy by pathogen effectors, NLR oligomeric complex undergoes structural modifications enabling the executor NLR to activate immune responses [13, 14, 18-20]. Currently, several NLR pairs have been functionally characterized. In *Arabidopsis thaliana*, two NLRs, Resistance to *Pseudomonas syringae* 4 (RPS4) with Resistance to *Ralstonia solanacearum* 1 (RRS1-R), cooperate genetically and molecularly to detect PopP2 and AvrRps4 effectors from root-infecting *Ralstonia solanacearum* and leaf-infecting *Pseudomonas syringae* bacteria, respectively. Molecular and structural analyses of RPS4/RRS1-R interactions showed that both receptors associate to form an inhibited, pre-activation receptor complex that is activated upon direct binding of effectors [15]. Two studies revealed that activation of RRS1-R/RPS4 occurs through the targeting of RRS1-R C-terminal WRKY domain by two unrelated effectors, PopP2 and AvrRps4 [18, 21]. PopP2, a member of the YopJ family effectors [22], was shown to acetylate key lysine residues in the invariant heptad of RRS1-R WRKY DNA-binding domain (ID). This acetylation promotes structural rearrangements that inhibit RRS1-R DNA-binding activity and disrupt RRS1-R intramolecular interactions [23] triggering activation of RPS4-dependent immunity. PopP2 uses the same lysine acetylation strategy to target multiple defense-promoting WRKY transcription factors. In the absence of RRS1-R/RPS4 recognition, PopP2 acetylation dislodges WRKY proteins from their DNA-binding sites and disables their *trans*-activating functions needed for defense gene expression [18]. In rice, the CNL pair RGA5/RGA4 cooperates genetically and physically in the recognition of two unrelated effectors of *Magnaporthe oryzae*, AVR-PIA and AVR1-CO39 [14, 24, 25]. The RGA5 NLR has an HMA domain, behaving like an ID that specifically binds both effectors. Interestingly, recognition of AVR-PIK effectors from *M. oryzae* is also mediated by an HMA ID contained in PIK-1 functioning with PIK-2 [19, 26].

These different examples reveal an original and sophisticated strategy used by plants to detect specific pathogen effectors by directly integrating an effector target decoy into immune receptors, triggering a specific immune reaction. This creates an effective surveillance mechanism for potent bacterial virulence activities which cannot be easily dispensed with by the pathogens. In this context, NLR-IDs can be considered as a very useful resource for the identification of yet uncovered (i) virulence targets of effectors and (ii) NLRs sensing functions of specific effectors. Additionally, the functional characterization of various IDs could be exploited to engineer NLR-IDs with either targeted or extended recognition capabilities for specific pathogen effectors.

In this study, we have cloned a subset of highly confident IDs from NLRs present in seven model and crop plant species. Our goal was to identify new ID/pathogen effector pairs highlighting potentially new effectors virulence targets. As a proof of concept, we screened in a yeast two-hybrid (Y2H) assay all possible pairwise interactions between the cloned IDs and a set of the 33 core type III effectors (T3Es) [27] from *Ralstonia solanacearum*, the model bacterial pathogen responsible of the bacterial wilt disease, affecting various host plant species [28]. In this screening, we used a library of cloned T3Es previously described [29]. This Y2H screen yielded ID interactors for 4 different T3Es: RipW, RipAE, RipS3 and PopP2. We tested the i*n planta* interaction for these different potential effector-ID pairs. Subsequently, we decided to focus our attention to a new WRKY ID containing NLR from Soybean (ID#85) interacting with PopP2.

This work aims at providing a resource for the community interested in screening plant pathogen effector repertoires to broaden the knowledge of virulence targets and provide a toolkit to design new NLR-IDs efficient for pathogen recognition.

## RESULTS

### A library of cloned NLR integrated domain (ID) from model plants

Our strategy was to clone IDs from NLRs of crop and non-crop model plants in order to obtain a curated library representing most of the known plant ID diversity. We chose a set of seven model plant species: *Arabidopsis thaliana* (*At*), *Solanum lycopersicum* (*Sl*), *Oryzae sativa* (*Os*), *Glycine Max* (*Gm*), *Medicago truncatula* (*Mt*), *Malus domestica* (*Md*) and *Brachypodium distachyon* (*Bd*). At first, *A. thaliana* and *B. distachyon* were selected as two model dicotyledones and monocotyledons species, respectively. Then, *S. lycopersicum* and *O. sativa* were added as they represent well studied crop species. The library was broadened by adding the two legume models: *M. truncatula* and *G. max*. The apple tree (*M. domestica)* was also included as this plant was reported to have a great number of NLR-IDs in its genome [10, 11].

To curate *NLR-ID* genes, we started with the gene list from two recent papers that scanned around 40 plant genomes for the presence of ID encoded in *NLR* gene sequences [10, 11]. For each of the 7 plant species that we selected, we manually curated *NLR-ID* genes by applying a specific pipeline to validate the gene-ID model. Each gene model was analyzed to confirm the integration of the ID within the considered *NLR* gene by identifying the relative transcripts (from available EST or RNaseq data). For Mt and Md, more recent genomic data [30, 31] than was analyzed previously, with more accurate gene models allowed us to re-evaluate the stringency of the *NLR-ID* identification. As a result, many early predictions were actually not annotated as NLR and many previously suspected ID were in fact colinear but distinct to the *NLR* genes. Finally, IDs sequences from *NLR-IDs* genes respecting these criteria were subsequent cloned following a specific procedure (Fig.S1). We decided to clone only the integrated domain to specifically address the decoy-effector interaction and prevent possible gene-expression issues in yeast.

Our stringent pipeline significantly reduced the number of predicted IDs in these seven plant genomes (from 252 originally predicted (Sarris *et al*., 2016) to 79 IDs, Fig.S2). We succeeded to clone 52 out of these 79 IDs, yielding a library containing 31 unique pfam domains (Table 1). In terms of diversity, our library represents 76 % of all pfam domains identified in the five species *At, Sl, Bd, Os* and *Gm* by Sarris and colleagues [11]. In *At*, 11 IDs have been cloned representing 8 unique ID with a high representation of BRX and WRKY domains (Table 1). Our library contains 19 *Sl*-ID mostly represented by DUF3542 described as a domain containing F-Box, Cupin_1 and DUF4377 domains (Table 1). We have cloned 9 *Os*-ID and 7 *Mt*-ID representing a good functional diversity with respectively 9 and 4 pfam domains (Table 1). We also added two *Gm*-ID (WRKY and Zf-Bed/ATG16), two *Md*-ID (Suc_Fer-like and Pkinase/Ef_hand_5). Finally, the library was enriched with 3 *Bd*-ID (Table1). We did not manage to clone all ID selected specifically from *Bd* (16 IDs selected to 3 IDs cloned) and *Gm* (11 IDs selected to 2 IDs cloned).

**Table 1:**
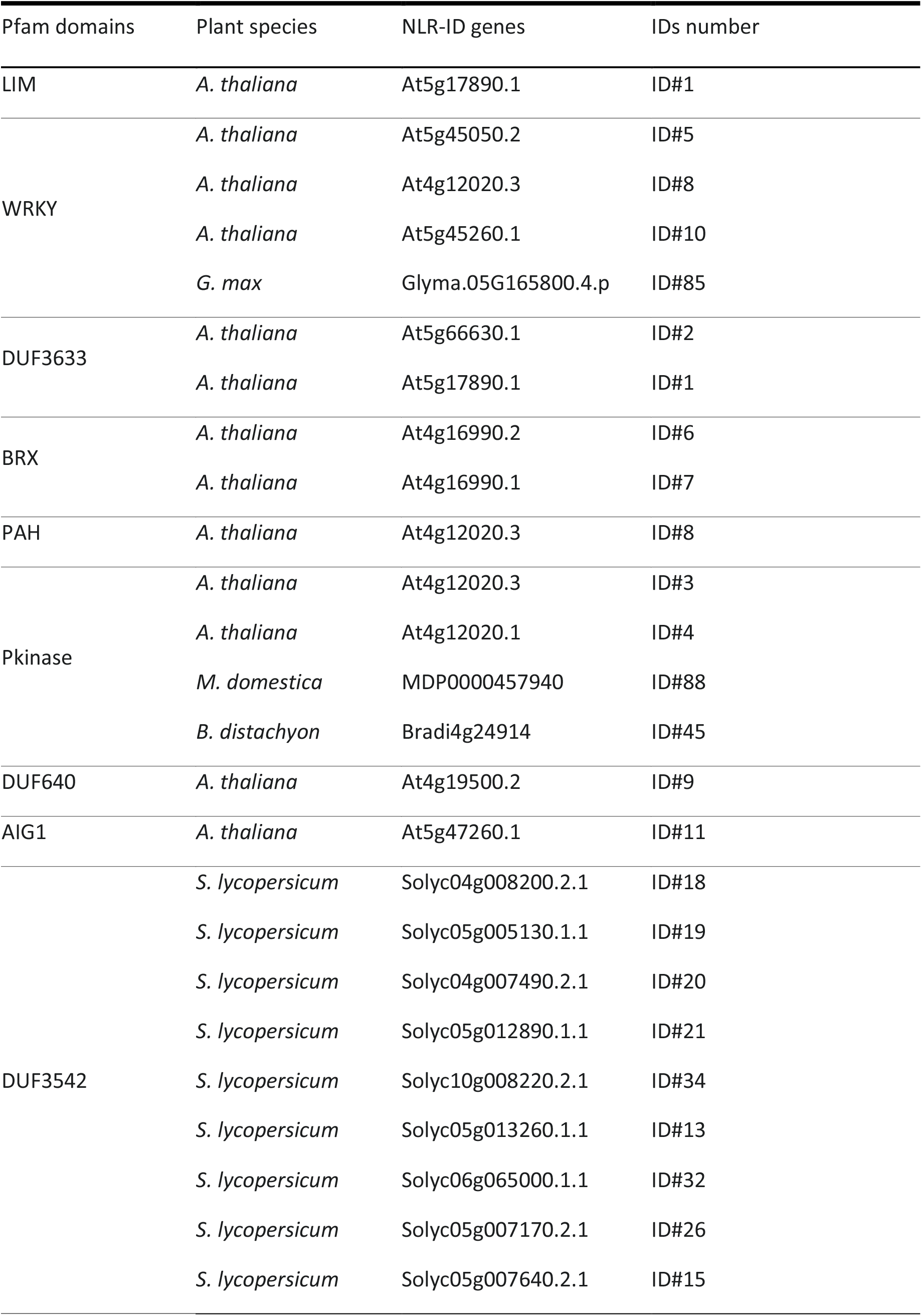
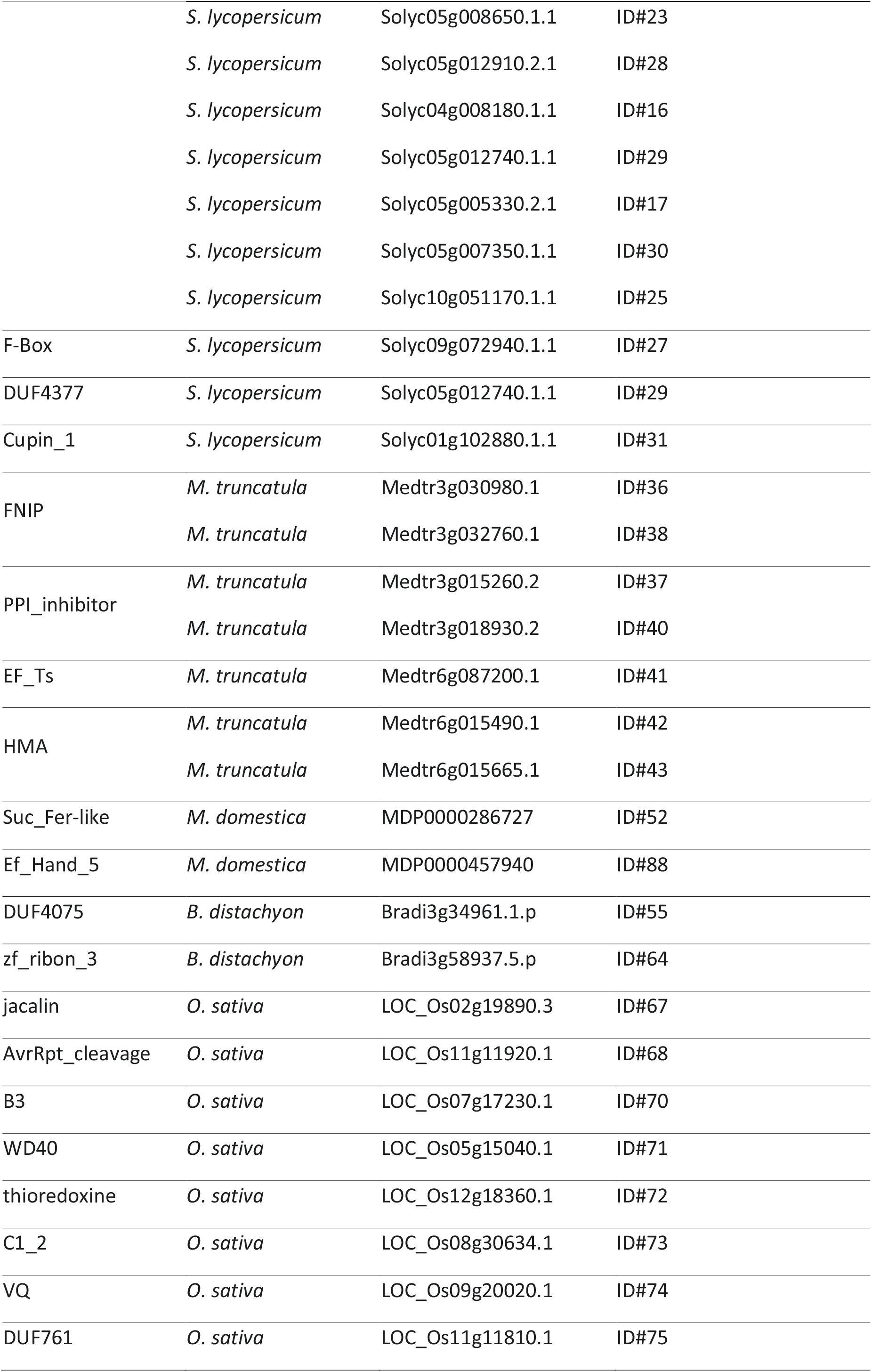
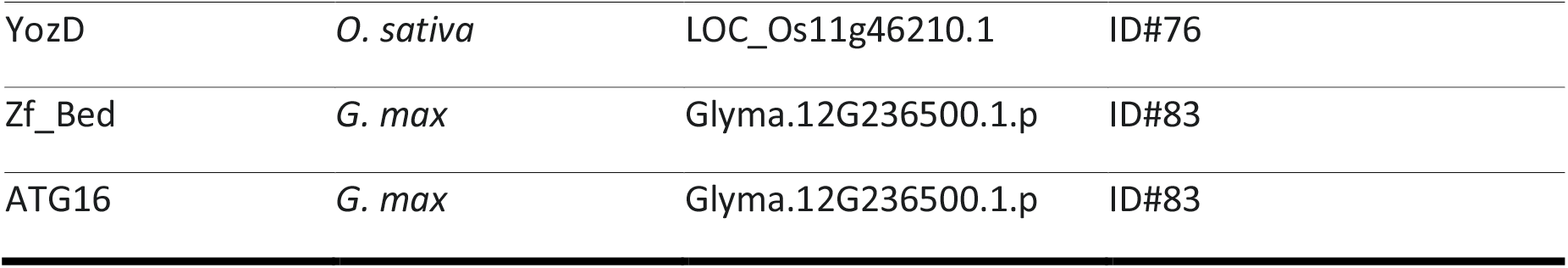
List of Integrated Decoy (ID) domains considered in this study (Pfam domain, species of origin and accession number).

### Yeast-two-Hybrid screening reveals IDs as potential virulence effector targets

Atypical domains integrated in NLR have been described to mimic true virulence effector targets. Consequently, using a Y2H assay, we investigated whether some *Ralstonia solanacearum* T3Es could interact with the cloned IDs. *R. solanacearum* is a species complex containing a large diversity of strains, several of which can provoke the bacterial wilt disease on a wide range of host plants. Plant pathogenic *R. solanacearum* strains share a large collection of type III effectors, directly targeting host processes for successful pathogen infection. We chose to screen the library with a collection of mostly conserved *R. solanacearum* T3Es [27]. The T3Es used in the screening are: RipA2, RipAB, RipAC, RipAD, RipAE, RipAI, RipAJ, RipAM, RipAN, RipAO, RipAY, RipB, RipC1, RipD, RipE1, RipG3, RipG5, RipG7, RipH1, RipH2, RipH3, RipM, RipN, RipP2(PopP2), RipR, RipS3, RipU, RipV1, RipW, RipX, RipY, RipZ. In Y2H assays, selected T3Es and cloned ID were used either as prey and bait proteins by fusing their C-terminal part with the GAL4 activation domain or the LEXA binding domain, respectively (see section Experimental Procedures). We identified and confirmed T3E/ID interaction for 17 yeast clones pairs growing on selective media (Fig.1, Fig.S3). Fig.S11 contains all the yeast-two-hybrid raw matrices from which Fig.1 and Fig.S3 were assembled. Two tomato DUF3542, ID#30 and ID#23, interacted respectively with RipS3, RipAE and RipW, RipAE (Fig.1A,B,E). The ID#3 clone that contains two At kinase domains, was found to interact with RipAE (Fig.1B). Subcloning of ID#3 into two distinct domains (ID#3A and ID#3B) revealed that only ID#3A retained its ability to interact with RipAE (Fig.1D). ID#29, a DUF3549, produced 11 unique interactions and was not further considered because of these too many interactions (Fig.S3). We also identified that ID#85, a WRKY domain from Soybean, interacted with PopP2 effector previously found as targeting many WRKY transcription factors *in planta* [18, 21]. All together, these interacting data highlight the potential of the yeast ID library to identify new potential effector targets. As show in Fig.S4, a few ID clones (ID#58, ID#64, ID#19) were not detected with the right size in yeast total protein extracts and couldn’t be considered as being screened.

**Figure 1.**
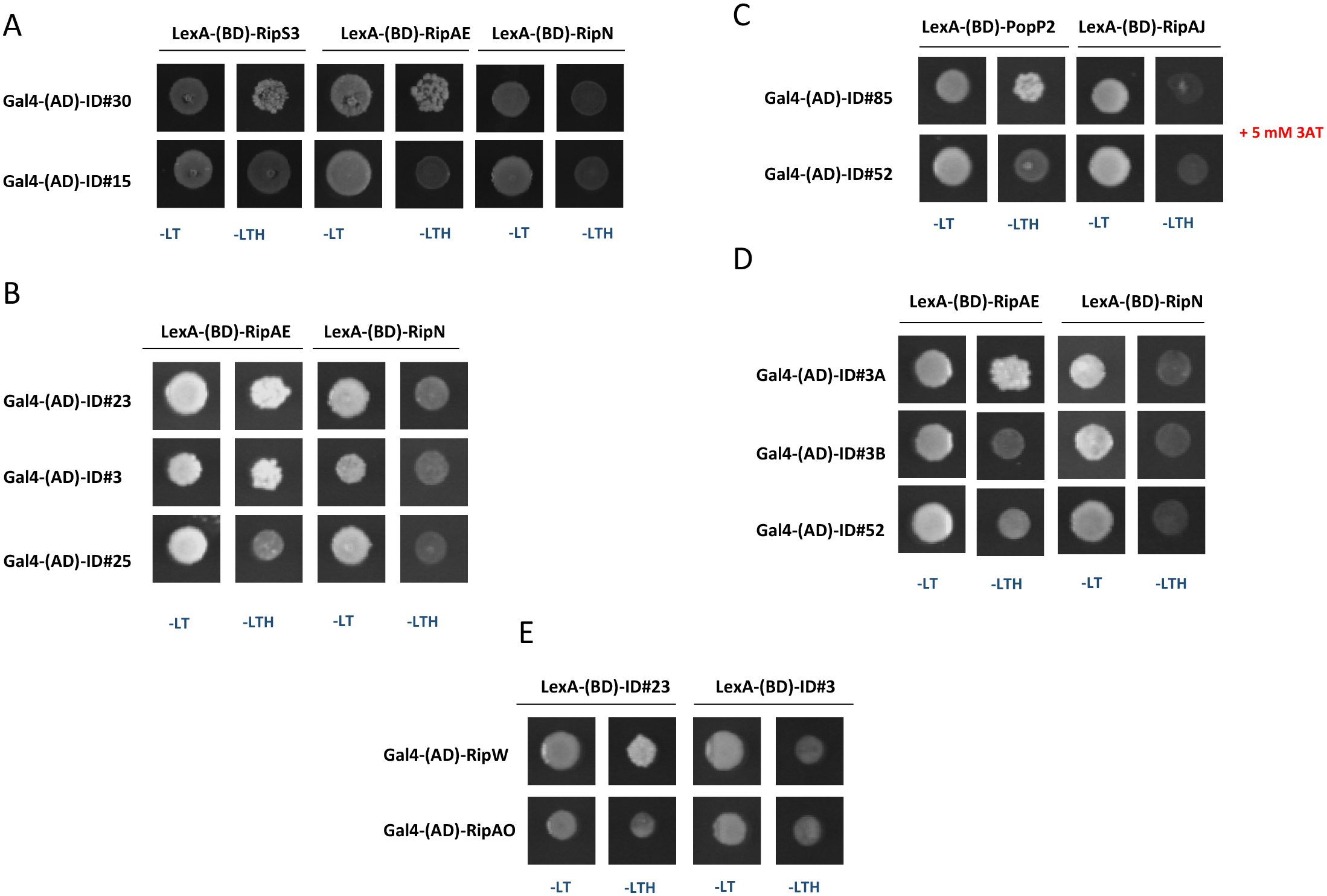
Yeast-two-hybrid screening reveals potential interactions between *R. solanacearum* T3Es and IDs. The interactions between cloned IDs and core T3Es were screened in a yeast-two-hybrid assay in both possible pairwise combinations. 17 independent yeast diploid clones were grown on selective media (SD-base minus leucine, tryptophan and histidine, -LTH) representing 17 different interactions (A, B, C, D, E and FigS3). The screening of LexA-(BD)-T3Es/Gal4-(AD)-IDs (A, B, C, D) led to the detection of the following interactions pairs: (A) RipS3/ID#30, RipAE/ID#30 with LexA-RipN and Gal4-ID#15 used as control; (B) RipAE/ID#23 and RipAE/ID#3 with LexA-(BD)-RipN and Gal4-(AD)-ID#25 used as negative control; (C) PopP2/ID#85 with LexA-RipAJ and Gal4-ID#52 used as negative control. For this assay, the media was supplemented with 3-amino-1,2,4-triazole (3AT) to prevent autoactivation of LexA-(BD)-PopP2 bait construct; (D) RipAE/ID#3A, RipAE/ID#3B with LexA-(BD)-RipN and Gal4-(AD)-ID#52 used as negative control. The constructs ID#3A and ID#3B have been generated by separating the two protein kinase domains of ID#3. Finally, the screening was also performed in the other orientation: LexA-(BD)-IDs/Gal4-(AD)-T3Es. From these experiments, only one combination led to growth in –LTH media (E) revealing the interaction between RipW and ID#3, with LexA-(BD)-ID#3 and Gal4-(AD)-RipAO used as negative control. For each experiments, the mating was checked by plating the yeast clones on non-selective media (SD base minus leucine and tryptophan, -LT). Both pairwise combinations were conducted in two independent replicates.

### ID-T3E pairs confirmed *in planta* interaction

For *in planta* validation of the interactions detected in yeast, a split luciferase assay was used by transiently expressing T3Es-Nluc (TE3s C-terminally fused with the amino terminal fragment of luciferase) and Cluc-IDs (IDs N-terminally fused with the carboxy-terminal fragment of luciferase) fusion proteins in *N. benthamiana* (Fig.2). An interaction between the two fusion proteins reconstitutes a functional luciferase enzyme, emitting light after addition of luciferin. This light-emission activity is detected and quantified with a luminometer. Thanks to this split luciferase assay, three of the interactions previously detected in yeast were validated *in planta*. Indeed, we showed that co-expression of Cluc-ID#3 with RipAE-Nluc produces a quantity of light significantly different from the Cluc-ID#3/RipG7-Nluc combination, used as negative control. In contrast, as a positive control, RipG7 and its plant target MSKA [32] fused with Nluc and Cluc, respectively, interacted when co-expressed *in planta*. The two kinase domains of ID#3, expressed as Cluc-ID#3A and Cluc-ID#3B fusion proteins, were also tested. ID#3A strongly interacts with RipAE-Nluc while a lower intensity of interaction was detected for ID#3B (Fig. 2), when no Y2H interaction could be detected for this latter pair (Fig.1D). RipAE-Nluc was also found to interact with Cluc-ID#30, confirming the interaction observed in yeast.

**Figure 2.**
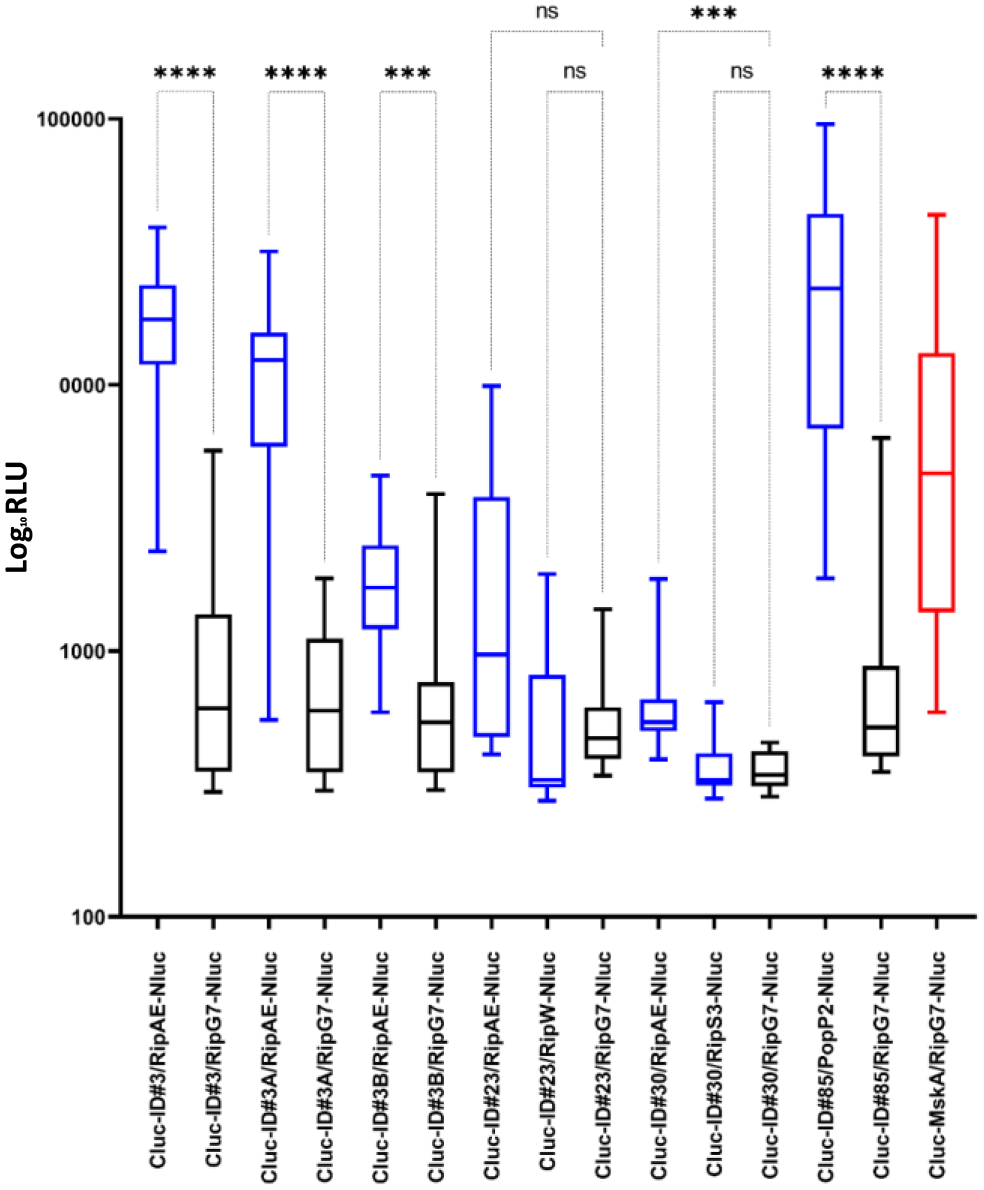
Validation of the in planta interaction between *R. solanacearum* T3Es and ID targets using a split-luciferase assay. A split luciferase assay was done by transient expression of Cluc-ID and T3E-Nluc in *N. benthamiana* leaves. The light emitted by the reconstitution of the luciferase was quantified by luminometer (Relative light unit, RLU). The experiment shows an interaction between RipAE and ID#3, ID#3A, ID#3B and ID#30. The PopP2 effector interacts *in planta* with ID#85 while the interaction between RipAE/RipS3 with ID#23 could not be confirmed. As negative controls, RipG7-Nluc was co-expressed with all Cluc-IDs (dark box-plot) while the co-expression of RipG7-Nluc with Cluc-MskA was used as positive control (red box-plot). This experiment was conducted in three independent biological replicates, containing each 16 technical replicates. All data points are represented in this figure. The non-parametric Kruskall-wallis statistical test was used; *** p-value < 0,0002; **** p-value < 0,0001.

Besides, ID#85 interacts with PopP2-Nluc and Cluc-ID30 interacts with RipAE-Nluc as the quantity of light was significantly higher from that observed with the negative control. Unfortunately, the other combinations tested (Cluc-ID#23/RipAE-Nluc, Cluc-ID#30/RipS3-Nluc and Cluc-ID#23/RipW-Nluc) were not validated *in planta* through this split luciferase assay.

### PopP2 relocalizes GmNLR-ID#85 in the plant nucleus wheres it does physically associate with its integrated WRKY domain

PopP2 was previously shown to target several WRKY transcription factors including the RRS1-R immune receptor that contains a WRKY decoy domain at its C-terminus [18, 21]. In our yeast screening, ID#85 was originated from *GmNLR-ID85*, a soybean *NLR* gene (Glyma.05g165800.4, predicted protein of 1351 residues) that integrates a WRKY domain at its C-terminus. ID#85 (residues 1105 to 1355 in GmNLR-ID85 protein) was found to interact with PopP2 both in yeast and *in planta* (Fig.1C, 2). To further validate this interaction *in planta*, we used a Förster resonance energy transfer (FRET) analyzed by fluorescence lifetime imaging microscopy (FLIM)-based approach to monitor protein interactions in living cells. First, we determined the subcellular localization of GmNLR-ID#85 C-terminally fused with Yellow fluorescent protein (YFP) and transiently expressed in *N. benthamiana* cells. Although the GmNLR-ID#85-YFP fusion protein expressed alone showed a nucleocytoplasmic distribution (Fig.3A), its co-expression with PopP2 fused with the Cyan fluorescent protein (PopP2-CFP) led to the accumulation of NLR-ID#85-YFP mostly exclusively in the nucleus (Fig.3A). Proper expression of the CFP and YFP fusion proteins was verified by immunoblot (Fig. 3B). This is reminiscent of RRS1-R nuclear accumulation shown to be increased in presence of PopP2 [33, 34]. Together, our observations suggest that PopP2 acts probably as cargo proteins and convey the GmNLR-ID#85 protein from the cytoplasm to the nucleus.

**Figure 3.**
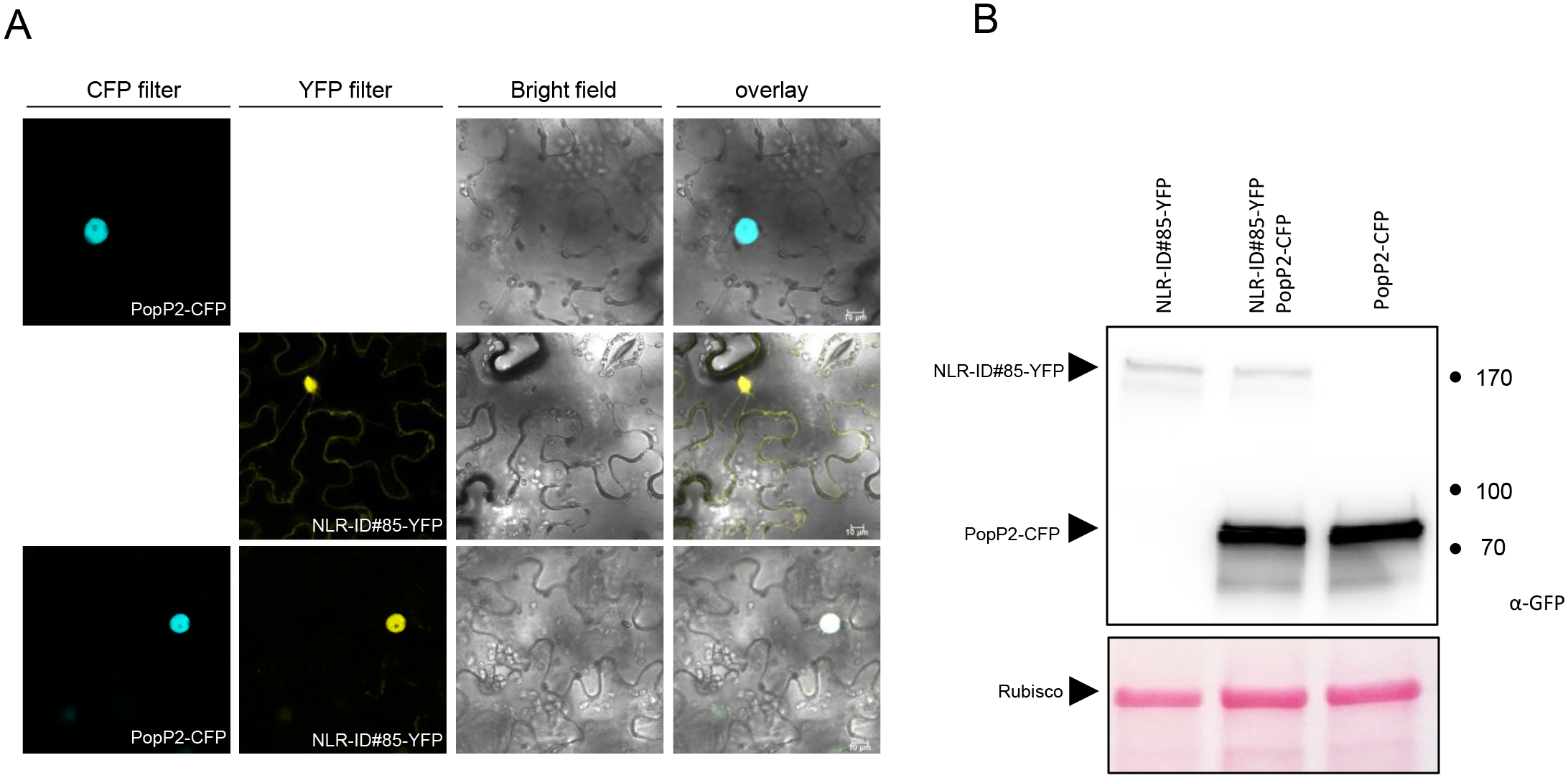
GmNLR-ID#85-YFP is relocalized to the plant nucleus in presence of PopP2-CFP. (A) The GmNLR-ID#85-YFP fusion protein was transiently expressed in *N. benthamiana* leaves with or without PopP2-CFP. The confocal fluorescence imaging shows that GmNLR-ID#85-YFP expressed alone shows a nucleocytoplasmic localization. Co-expression of GmNLR-ID#85-YFP with PopP2-CFP generates a nucleus-restricted YFP signal. (B) Expression of GmNLR-ID#85-YFP and PopP2-CFP fusion protein in *N. benthamiana* was checked by immunoblot with an anti-GFP antibody.

Although the co-localization data were unequivocal, the accumulation level of GmNLR-ID#85-YFP was too low to serve as FRET acceptor in combination with PopP2-CFP used as a donor. Unfortunately, the original ID#85 clone (aa 1105 to 1355) fused with YFP could also not accumulate at sufficient level *in planta* (data not shown). Therefore, we decided to re-clone a shorter version of the ID#85 (from residues 1246 to 1355), as previously done [18]. This domain, hereafter designated as WRKY-ID#85, was then fused with CFP to serve as FRET donor in FLIM-FRET measurements (Fig.4, Table 2). WRKY-ID#85 construct was designed to be similar in length to the RRS1-R WRKY domain previously shown be targeted by PopP2 [18]. When expressed alone or in presence of YFP, WRKY-ID#85-CFP displayed an average CFP lifetime of 2,86 ns ±0,014 and 2,79 ns ±0,017, respectively (Table 2). By contrast, co-expression of WRKY-ID#85-CFP with PopP2-YFP led to a significant decrease of the CFP lifetime (2.53 ns ±0,013) resulting in 11% of FRET efficiency, indicating that WRKY-ID#85 and PopP2 physically interact in the plant nucleus. This interaction was further validated by an independent GST pulldown assay in which 3HA-tagged WRKY-ID#85 protein transiently expressed in *N. benthamiana* was able to form a complex with GST-PopP2-6His but not with a GST-GUS-6His fusion protein used as a negative control (Fig.S6).

**Figure 4.**
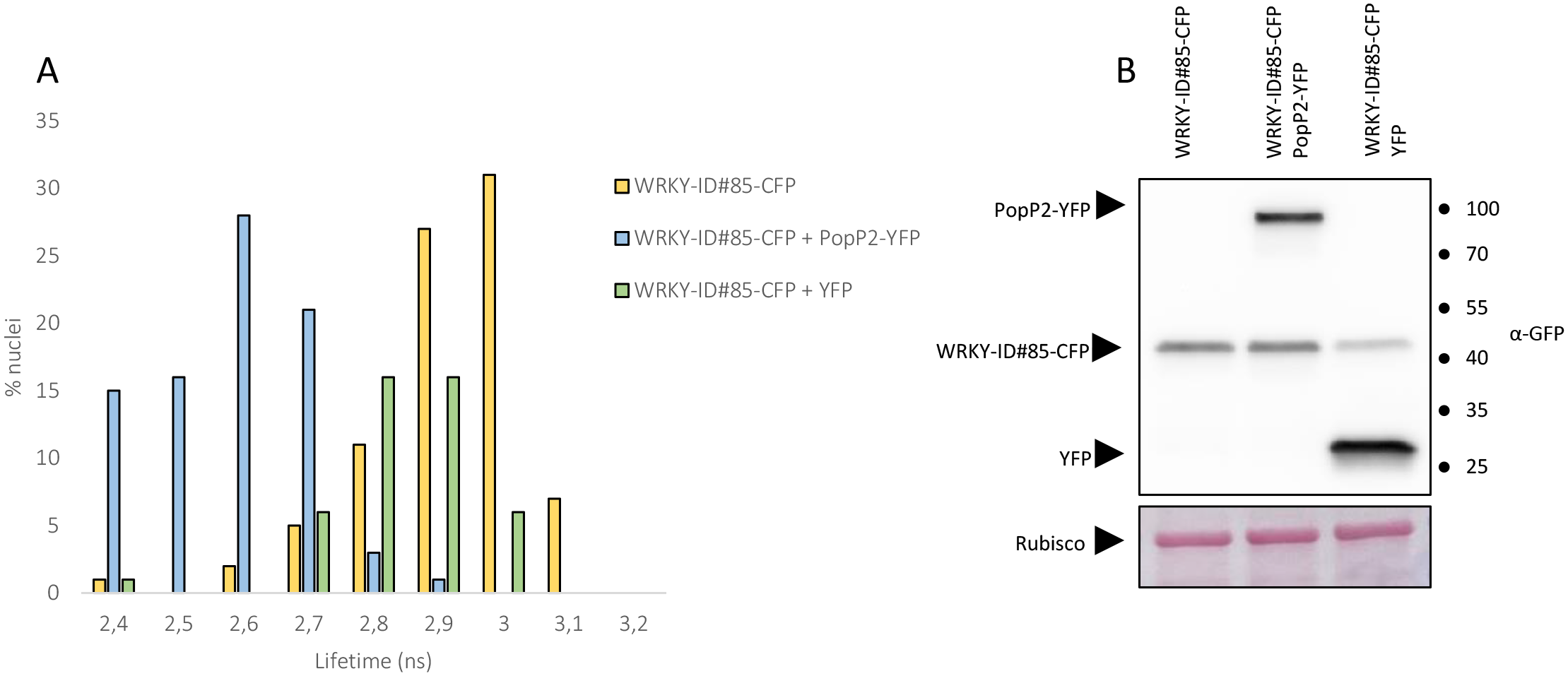
The WRKY domain of GmNLR-ID#85 physically interacts with PopP2 in the nucleus of *N. benthamiana* cells. The WRKY domain of GmNLR-ID#85 fused with CFP (WRKY-ID#85-CFP) was transiently expressed in *N. benthamiana* leaves either alone or with YFP or PopP2-YFP and CFP fluorescence lifetime was measured in relevant individual nuclei. (A) Histograms show the distribution of nuclei (%) according to CFP lifetime classes of WRKY-ID#85-CFP expressed alone (yellow bars) or in presence of PopP2-CFP (blue bars) or YFP (green bars). in the absence. (B) Accumulation of WRKY-ID#85-CFP, PopP2-YFP and YFP proteins in *N. benthamiana* cells was checked by immunoblot with an anti-GFP antibody.

**Table 2:** FLIM-FRET measurements showing a physical interaction betweenWRKY-ID#85 and PopP2 in the nucleus of *N. benthamiana* cells.

| Donor | Acceptor | Lifetime*<br>(ns) | SEM<sup>1</sup> | N<sup>2</sup> | E (%)<sup>3</sup> | p-value<sup>4</sup> |
| --- | --- | --- | --- | --- | --- | --- |
| WRKY-ID#85-CFP | - | 2,86 | 0,014 | 84 |  |  |
| WRKY-ID#85-CFP | PopP2-YFP | 2,53 | 0,013 | 84 | 11,57 | 7,61E-39 |
| WRKY-ID#85-CFP | YFP | 2,79 | 0,017 | 45 | 2,47 | 2,68E-03 |

\*Mean lifetime in nanosecond (ns). For each nucleus, average fluorescence decay profiles were plotted and fitted with exponential function using a non linear square estimation procedure and the mean lifetime was calculated according to $\tau = \sum \alpha_i t_i^2 / \sum \alpha_i t_i$ with $I(t) = \sum \alpha_i e^{-t/t_i}$ , <sup>1</sup>Standard error of the mean, <sup>2</sup>Total number of measured nuclei, <sup>3</sup>E (%): Energy transfer efficiency: $E=1 - (\tau_{DA}/\tau_D)$ , and <sup>4</sup>p-value of the difference between the donor lifetimes in the absence and in the presence of acceptor (Student's *t* test).

### The atypical WRKY domain of GmNLR-ID#85 does not behave as a substrate of PopP2 acetyltransferase

Previously published data showed that PopP2 acetylates two lysine residues located in the highly conserved core WRKYGQK heptad of WRKY transcription factors, including AtWRKY52 that corresponds to the RRS1-R immune receptor [18, 21]. Protein alignment of WRKY-ID#85 with the WRKY domain of RRS1-R revealed that the heptad sequence present in GmNLR-ID#85 diverges from that of RRS1-R in its last two residues, W^1277^RKYGKR^1283^ and W^1215^RKYGQK^1221^, respectively (Fig.S7). We thus determined whether PopP2 could acetylate this atypical ID#85 WRKY domain. For this, a GST-WRKY-ID#85-His6 fusion protein was co-expressed with His6 epitope-tagged PopP2 (His6-PopP2) or PopP2^C321A^ (His6-PopP2^C321A^). GST-ID#85-His6 from both bacterial extracts was affinity purified on glutathione sepharose. The RRS1-R WRKY domain contained in the C-terminal portion of RRS1-R (position 1190-1379, hereinafter called RRS1-RCterm) was previously described as behaving as a direct substrate of PopP2 acetyltransferase (Le Roux et al., 2015). We therefore included as positive control a GST-RRS1-R_Cterm_-His6 fusion protein (Fig. S8A). Immunoblot analysis of GST-WRKY-ID#85-His6 co-expressed with PopP2 or PopP2^C321A^ led to the detection of a low intensity signal with an anti-AcK antibody, suggesting that GST-WRKY-ID#85-His6 is acetylated in *E. coli* independently of the presence of enzymatically active PopP2 (Fig.S8A). To circumvent this problem, acetylation of WRKY-ID#85 by PopP2 was investigated in *N. benthamiana* (Fig.S8B). WRKY-ID#85 and RRS1-R_Cterm_ were C-terminally fused to a YFP or a eGFP tag (WRKY-ID#85-YFP and RRS1-R_Cterm_-eGFP, respectively) and co-expressed either with triple hemagglutinin (HA) epitope-tagged wild-type PopP2 or catalytic mutant PopP2^C321A^. Both WRKY-ID#85-YFP and RRS1-R_Cterm_-eGFP and were purified from protein extracts using a GFP affinity matrix. Contrary to what was observed with RRS1-R_Cterm_-eGFP, acetylated forms of WRKY-ID#85-YFP expressed with wild-type PopP2 could not be detected with an anti-AcK antibody (Fig. S7B), suggesting that PopP2 is unable to cause acetylation of Lys1279 and Lys1282 residues located in the divergent heptad of WRKY-ID#85. Together, our data indicate that although WRKY-ID#85 physically interacts with PopP2, it does not behave as a substrate of PopP2 acetyltransferase activity.

### The divergent heptad of WRKY ID#85 maintains autoinhibition of RRS1-R

Previous studies have shown that RRS1-R WRKY domain negatively regulates the RPS4-RRS1-R complex [23]. Importantly, the last Lys residue of RRS1-R WRKY heptad in position 1221 is critical for NLR activation since its substitution either with Q or R residues (K1221Q and K1221R) renders RRS1-R autoactive and nonautoactive, respectively (Le Roux et al., 2015). To investigate the functional properties of the divergent heptad of WRKY ID#85 in NLR activation, its last two residues were introduced into the conserved WRKY heptad of RRS1-R (the native WRKYG<u>Q^1220^K^1221^ heptad sequence of RRS1-R was replaced with</u> WRKYG<u>K^1220^R^1221^)</u>, resulting in the RRS1-R^WRKYGKR^ variant. Stable T2 transgenic lines (lines #25 to #28) expressing RPS4 and RRS1-R^WRKYGKR^ isoforms under their genomic 5’ and 3’ regulatory sequences were made in *rps4-21 rrs1-1* (Ws-2 background, hereinafter called *r4r1*) and root-inoculated with *R. solanacearum* GMI1000 strain to assess for PopP2-triggered activation of RPS4/RRS1-R-dependent immunity. Control lines expressing both wild-type *RPS4* and *RRS1*-R (RPS4/RRS1-R^Ws-2^) were also generated in the same genetic background (*r4r1* lines #9 to #12) and included in our root pathogen assay (Fig.5, Fig.S9). While lines #9-12 showed a resistance phenotype in response to GMI1000 reflecting proper activation of the RPS4/RRS1-R pair by PopP2, transgenics (lines #25-28) expressing RRS1-R^WRKYGKR^ developed wilting symptoms comparable to those of the untransformed *r4r1* mutant. These data indicate that PopP2, although targeting the divergent heptad of WRKY ID#85, is unable to relieve the autoinhibitory intramolecular interactions occurring in RRS1-R^WRKYGKR^.

**Figure 5.**
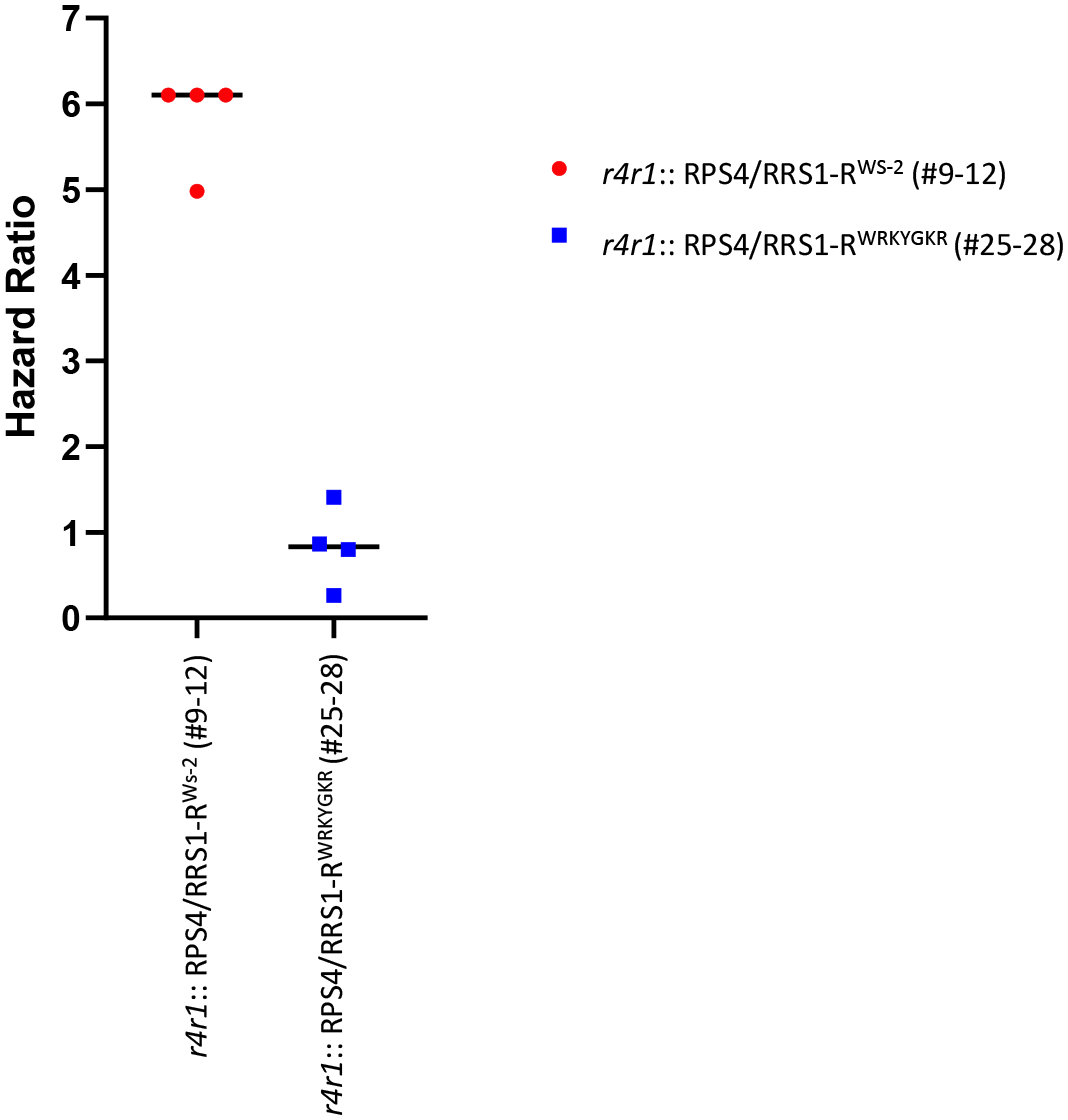
Introduction of the WRKY-ID#85 divergent heptad in RRS1-R immune receptor compromises its responsiveness to PopP2. The hazard ratios obtained for 8 individual logrank test comparisons (Figure S9) are plotted here, with their associated median. Each comparison was made with the reference *A. thaliana rps4-21 rrs1-1* double mutant (noted *r4r1)*. In red, 4 individual T2 transgenic lines expressing RPS4/RRS1-R (Wild-type sequence from Ws-2 ecotype, noted *r4r1*::RPS4/RRS1-R^Ws-2^) in the double mutant background. All 4 lines showed a resistance phenotype upon root infection with the GMI1000 strain (three lines showing no symptoms at all, see Figure S8), unlike the susceptible *r4r1* double mutant (Hazard ratio media=6.1). In blue, 4 individual T2 transgenic lines expressing RPS4/RRS1-R^WRKYGKR^ (introduction of the WRKY-ID#85 divergent heptad in RRS1-R) in the double mutant background (noted *r4r1*::RPS4/RRS1-R^WRKYGKR^). Here the hazard ratio close to 1 (median=0,83) indicates that the wilting of the double mutant line in response to GMI1000 is similar to that of the RPS4/RRS1-R^WRKYGKR^ containing lines. A Mann Whitney test (P-value=0,0286) shows that both hazard ratio groups are significantly different. Individual line comparison P-values are given for the logrank test in Figure S9.

## DISCUSSION

In this work, we aimed at cloning several IDs contained in NLR from a set of model and crop plants. As we wanted to generate a screenable Y2H library of ID, we first needed to curate the NLR gene models to have a high degree of certainty that the cloned IDs were indeed genuine IDs. To this goal, we only cloned ID for which we could find evidence of gene expression by identifying ESTs or RNAseq traces for the cognate gene models. Furthermore, for two plants species, the availability of more recent (and more accurate) whole genome sequences, drastically changed the prediction of IDs. This was the case for Apple (*Malus domestica*) and *Medicago truncatula* were original prediction identified 93 and 48 IDs, respectively [11]. While our curation of newly available genomes [30, 31] only identified 2 and 9 potential NLR-IDs in these two species. Our curation also reduced by approximately half the number of credible ID in Rice and Soybean (from 22 to 10 in Rice and 25 to 11 in Soybean).

We succeeded to clone 52 individual ID out of the 79 that we originally set out to clone. We could not explain why both *Brachypodium distachyon* and Soybean (*Glycine max*) seemed recalcitrant as we only managed to clone 3/16 and 2/11 of the predicted ID in these two species. All cloning were done from cDNA prepared from total RNAs, we can imagine that some very low expression level could be the issue for some of the failed cloning. Although this should not be specific to a given plant species.

Without considering the bias introduced by the previous over prediction of NLR-IDs, our library of 52 ID clones, contains 31 unique Pfam domains representing 76% of the Pfam domain originally predicted for these 7 plant species.

Our original aim was to provide means to screen the ID library for interactions with plant pathogen effectors. The underlying idea being that these IDs have been selected and maintained through evolution of *NLR* genes in these plants and therefore most likely behave as molecular decoys for the specific recognition of yet unidentified virulence effectors. These atypical domains in NLRs can guide us to identify the actual *bona fide* targets of some virulence effectors and provide means to engineer new resistance genes. This strategy was outline previously [35].

We exploited our knowledge [27, 36] and availability of a set of conserved *Ralstonia solanacearum* T3Es [6] cloned and readily available [29]. We used a Yeast-2-Hybrid mating strategy, as a quick and efficient method to screen for interactions between the collection of 31 cloned *R. solanacearum* T3Es and the library of 52 IDs clones. Although heterologous, Y2H has been the method of choice for identifying host targets of diverse pathogen effectors in many dedicated or large scale studies [29]. This allowed us to identify several candidate T3E-ID pairs involving five different IDs (ID#3, #23, #29, #30 and #85) and four different *R. solanacearum* T3Es (RipAE, PopP2, RipS3 and RipW). In order to further validate these putative protein-protein interactions, we managed to confirm the *in planta* interaction using a split-Luciferase system of the following T3E-ID pairs: #ID3A (kinase domain from *Arabidopsis*) with RipAE; #ID30 (DUF3542 from tomato) with RipAE and #ID85 (WRKY domain from Soybean) with PopP2. Interestingly, both kinases (#ID3A and #ID3B) present in tandem in the #ID3 domain cloned from Arabidopsis (At4g12020.3) could interact *in planta* with RipAE, when only the #ID3A showed an interaction with RipAE in Y2H. The other Y2H interactors that could not be confirmed *in planta*, should not be ruled out, as every protein-protein interaction method has its own false negative bias.

To further prove the interest of screening this ID library resource, we followed up on the ID#85 (WRKY domain integrated in GmNLR-ID#85, an *NLR* gene from Soybean) and PopP2, reminiscent of the RRS1-R WRKY domain acting as a molecular decoy that mimics the true targets of PopP2, the defensive WRKY transcription factors [18, 21]. We showed that the NLR-ID#85 from soybean is targeted to the nucleus of epidermal *N. benthamiana* cells when co-expressed with PopP2. Furthermore WRKY-ID#85 interacts *in planta* with PopP2 as was evidenced by split-Luciferase assay and FLIM-FRET analysis.

We observed that the WRKY domain of GmNLR-ID#85-ID#85 was atypical in its amino-acid sequence; with the “WRKYGKR” heptade rather than the more canonical “WRKYGQK” (Fig.S7). Unlike RRS1-R WRKY domain, acetylated forms of WRKY-ID#85 couldn’t be immunodetected in presence of wild-type PopP2, nor in a *E. coli* heterologous system, nor in an *in planta* assay. Furthermore, integration of this atypical WRKY heptad in RRS1-R resulted in loss of responsiveness to PopP2 delivered upon root inoculation with the *R. solanacearum* strain GMI1000. This is consistent with the apparent inability of PopP2 to acetylate this divergent WRKY heptad.

We observed that orthologs of GmNLR-ID#85 within the Fabaceae family could also harbor an ID-WRKY in the C-terminal part of the protein (Fig.S10). Interestingly, the sequence of the WRKY heptade can be canonical (“WRKYGQK”, *A. hypogea* XP_025625112.2, *V. unguiculata* QCD84554.1, *V. radiata* XP_014510449.1) or divergent (“WRKYGKK” *V. angularis* XP_01743563.1; “WRKYGKR” *G. max* KRH59109.1, GmNLR-ID#85 in this work). This could indicate that these different WRKY integrated domains could represent different stages of evolved IDs [37] some being active in immune signaling (for the canonical-WRKY containing NLRs) and some being under less stringent selection and potentially having lost some of their original features. This could be the case for GmNLR-ID#85. Alternatively, such an ID in GmNLR-ID#85 could sense pathogen interference with WRKY transcription factors involving a different mechanism than that used by PopP2 acetyltransferase. At this stage we cannot rule out that the contribution to immunity of GmNLR-ID#85 is different from that of RRS1-R, hence the absence of complementation in *Arabidopsis*.

We believe that though screening for interaction between our library of IDs and T3E repertoire we have uncovered a similar but different NLR potentially serving as a decoy for the PopP2 effector from *R. solanacearum*. Other interesting ID-T3E pairs could be explored further, namely the *in planta* confirmed interactions, #ID3A (kinase domain from *Arabidopsis*) with RipAE and #ID30 (DUF3542 from tomato) also with RipAE.

Our 52 cloned ID library could be further completed by adding new IDs from other plant species to increase the Pfam diversity coverage. We could imagine also adding more IDs from other ecotypes of the plants already sampled to increase the natural diversity of available ID sequences. This was done recently for the Heavy-Metal Associated (HMA) IDs [38]. At this stage, this library is already a great representation of the known diversity of IDs and is available for screening with other pathogen effector repertoires. This useful resource could prove important in unraveling both the virulence and immune-triggering functions of T3Es, as well as providing leads for future *NLR* gene engineering.

### EXPERIMENTAL PROCEDURES

#### Plant material and culture conditions

*Nicotiana benthamiana* plants were cultivated in a growth chamber at 19-21°C under 16h light/8h dark photoperiod. Arabidopsis plants (Ws-2 and *rps4-21 rrs1-1* double mutant (in Ws-2 genetic background)) were grown in Jiffy pots under controlled conditions (22°C, 60% relative humidity, 125 µE/M2/s fluorescent illumination, 8 h light:16h dark cycle).

#### ID cloning

IDs were cloned by following a specific strategy applied as shown in FigS1: (1) an isolated ID was cloned with 501 upstream and 501 downstream flanking nucleotides; (2) When the ID is surrounded with other canonical domains (RPW8, TIR/CC, NB-ARC, LRR), it was cloned at the border of these conserved-domains; and (3) if the ID is followed by another ID, both IDs were cloned together following the previous rules. All IDs were amplified from cDNAs used as PCR templates. Total RNAs were extracted from relevant plant tissues, and cDNAs were synthesized using classical procedures.

#### Bacterial strains, Yeast strains and Growth conditions

*Escherichia coli* DH5α, Rosetta (DE3), and *Agrobacterium tumefaciens* GV3101, GV3103 or C58C1 cells were grown in Luria-Bertani (LB) medium at 37 and 28°C, respectively. For liquid culture, *A. tumefaciens* cells were grown in yeast extract beef (YEB) liquid medium at 28°C, overnight. Antibiotics were used at this following final concentrations (μg/mL): for *E. coli*: kanamycin (50), tetracycline (5), gentamicin (10), chloramphenicol (25), carbenicillin (50) and spectinomycin (50); for *A. tumefaciens*: kanamycin (25), tetracycline (5), gentamicin (20) and carbenicillin (25).

For Y2H experiments, two different yeast strains, L40 and Y187, were used for mating. The Y2H plasmids pP6 [39] and pB27 [40] plasmids provided by Hybrigenics (Paris) were modified into pNP377 and pNP378 Gateway Destination vectors by inserting the chloramphenicol/ccdB resistance Gateway cassette (Invitrogen). Recombined pDEST pNP377 and pNP378 plasmids were introduced in yeast cells using the LiAc/SS carrier DNA/PEG method described here (see Yeast Transformation). Yeast cells were grown at 30°C on rich medium YPGA or yeast minimal media SD-base (Takara Bio) supplemented with histidine 20 mg/L, tryptophan 20 mg/L or leucine 10 mg/L. Interaction test by mating was performed following standard procedures.

#### Plasmids and Cloning

The IDs were amplified with Primestar Max DNA polymerase (Takara Bio) using as template either cDNA or genomic DNA prepared from several plant species: *Arabidopsis thaliana* Col-0, *Medicago truncatula* A17, *Brachypodium distachyon* Bd21-3, *Malus domestica* Golden delicious, *Oryzae sativa* nipponbare, *Solanum lycopersium* M82 and *Glycine max* Wm82. The PCR products were recombined into pDONR207 by Gateway technology (BP reaction) or were cloned by TOPO cloning into pENTRY-SD-TOPO. All clones were verified by sequencing. The genes of interest were then recombined in pNP377 (Gal4(AD)-GWY) and pNP378 (LexA(BD)-GWY) Destination vectors using LR clonase (Invitrogen) for subsequent yeast-two-hybrid experiments. For Agrobacterium-mediated transient expression in *N. benthamiana*, the following pDEST vectors were used: pAM-PAT-35S-GWY-CFP/YFP, pAM-PAT-35S-GWY-3HA or pBIN-35S-GWY-CFP/YFP. The genes of interest were recombined in these vectors using LR clonase (Invitrogen). For acetylation assays in *E. coli* cells, two pDEST vectors were used: pCDF-GST-GWY-6His and pDUET-6His-GWY. For Split-luciferase assay, the pDEST-Cluc-GWY and pDEST-GWY-Nluc vectors were used [41].

#### *Agrobacterium*-mediated transient expression in *N. benthamiana*

Agrobacterium cells were cultivated overnight in YEB media with appropriate antibiotics and were resuspended in infiltration medium (10 mM MES pH5.6, 10mM MgCl_2_ and 150 µM acetosyringone) to a final concentration of 2.5×10^8^ cfu/mL (OD_600_=0.25). For co-expression, bacterial suspensions were mixed in a 1:1 ratio. For split luciferase assay, OD_600_=0.125 of each bacterial suspension was used. After incubation at room temperature for 1 h in infiltration medium, bacteria were infiltrated into *N. benthamiana* leaves using a needle-less syringe. The infiltrated plants were incubated in growth chambers under controlled conditions. After 48 h, 4 leaf disks (8 mm of diameter) were harvested and proteins were extracted for immunoblot analysis.

#### Microscopy and FLIM-FRET analysis

CFP and YFP fluorescence was analyzed with a confocal laser scanning microscope (TCS SP8; Leica) using a x25 water immersion objective lens (numerical aperture 0.95; HCX PL APO CS2). CFP and YFP fluorescence was excited with the 458/514 nm ray line of the argon laser and recorded in one of the confocal channels in the 465-520/ 525-600nm emission range respectively. The images were acquired in the sequential mode using Leica LAS X software (version 3.0). Fluorescence lifetime measurements were performed in time domain using a streak camera. The light source is a 439 nm pulsed laser diode (PLP-10, Hamamatsu, Japan) delivering ultrafast picosecond pulses of light at a fundamental frequency of 2 MHz. All images were acquired with a 60x oil immersion lens (plan APO 1.4 N.A., IR) mounted on an inverted microscope (Eclipse TE2000E, Nikon, Japan). The fluorescence emission is directed back into the detection unit through a short pass filter and a band pass filter (483/32 nm). The detector is a streak camera (Streakscope C10627, Hamamatsu Photonics, Japan) coupled to a fast and high-sensitivity CCD camera (model C8800-53C, Hamamatsu, Japan). For each acquisition, average fluorescence decay profiles were plotted and lifetimes were estimated by fitting data with exponential function using a non-linear least-squares estimation procedure [42]. Fluorescence lifetime of the donor was experimentally measured in the presence and absence of the acceptor. FRET efficiency (E) was calculated by comparing the lifetime of the donor (#ID85-CFP) in the presence (τ_DA_) or absence (τ_D_) of the acceptor (PopP2-YFP): E=1-(τ_DA_)/(τ_D_). Statistical comparisons between control (donor) and assay (donor + acceptor) lifetime values were performed by Student *t* test.

#### Yeast transformation

Yeast cells were cultivated overnight on liquid YPGA medium and were used to inoculate a daily culture at OD_600_= 0.15. After 6 hours at 30°C, yeast cells were pelleted (3500 rpm, 5 min), washed in sterile water, and resuspended in 1 mL of LiAc 100 mM. For transformation, cells were with PEG 29%, LiAc 87 mM, Salmon sperm 240 ug/mL (previously denatured by boiling) and 100 ng of Y2H plasmid (pNP377 or pNP378 based). Yeast cells were then incubated at 30°C during 30 min and 42°C during 30 min. Transformed yeast cells were pelleted (7000 rpm, 15 sec) and resuspended in 200 µL of sterile water. Cells were plated in minimal SD media (SD-Leu or SD-Trp, for cells transformed with Gal4-(AD) and LexA-(BD) plasmids, respectively).

#### Yeast two Hybrid matrix

For mating, transformed L40 and Y187 yeast cells were harvested on plate and resuspended in liquid YPGA medium. Y2H experiments were performed in sterile 96-well plate by mixing yeast cells (ratio 0.2:0.2) in each well. The mating was performed overnight at 30°C under 180 rpm agitation. Diploïd yeast cells were plated in selective media SD–LT (to estimate mating efficiency) and SD–LTH (for selection of positive interactions). To prevent bait autoactivation, selective media were supplemented with 5 mM of 3-Amino-1,2,4-triazole (3-AT).

#### Bacterial and *in planta* acetylation assays

The acetylation assays were performed either in *E. coli* Rosetta (DE3) cells or *in planta* (*N. benthamiana*). Recombinant 6His-tagged proteins were produced in Rosetta (DE3) cells grown until OD_600_ 0.4 to 0.6 and induced with 250µM IPTG for 4 hours at 28°C. Bacterial cells were disrupted in protein extraction buffer I (50 mM Tris-HCl pH 7,5, 150 mM NaCl, 10 mM Sodium Butyrate (NaB, SIGMA), 0.1 % Triton-X-100, 6 M urea, 20 mM imidazole, 1mM PMSF) using a French Press cell disruptor. Resulting lysates were incubated for 30 min with Ni-NTA agarose beads (Qiagen) at 4°C. After incubation, beads were washed 3 times in protein extraction buffer I and Ni-NTA-bound proteins were denaturated in Laemmli 2X for 5 min at 95°C and subjected to immunoblot analysis. For the acetylation assays performed in *N. benthamiana*, 4 leaf disks (8 mm of diameter) were harvested and ground in liquid nitrogen. The total proteins were extracted with 1 mL of protein extraction buffer II (50 mM Tris-HCl pH7.5, 150 mM NaCl, 10 mM EDTA, 0.2% triton-X-100, 2mM DTT, 10 mM NaB and 1X plant protease inhibitor cocktail (SIGMA)). GFP-tagged proteins were immunoprecipitated with GFP-beads (Chromotek) during 1 hours at 4°C. After incubation, GFP beads were washed 3 times in protein extraction buffer II and purified protein were denaturated in Laemmli 2X for 5 min at 95°C. Protein samples were then subjected to immunoblot analysis.

#### *In vitro* GST pull-down

Recombinant GST-PopP2-6His and GST-GUS-6His fusion proteins were produced in transformed *E. coli* Rosetta (DE3) cells (Novagen) (see bacterial acetylation assay). Bacterial cells were disrupted in PBS protein extraction buffer (1X PBS, 0.5% Triton-X-100, 1 mM PMSF and 1X complete EDTA-Free protease inhibitor cocktail (Sigma)) using a French Press cell disruptor. Resulting lysates were incubated with Glutathione Sepharose 4B beads (GE Healthcare) for 2 hours at 4°C. After incubation, beads were washed 3 times in PBS protein extraction buffer. Then, the bound proteins were incubated for 2 hours at 4°C with an equal volume of a plant protein extract containing the WRKY-ID#85-3HA protein. Briefly, this plant protein extract was prepared from 3g of fresh *N. benthamiana* leaves transiently overexpressing WRKY-ID#85-3HA. Total proteins were resuspended in PBS protein extraction buffer. Glutathione beads were then washed three times in x mL of PBS extraction buffer. The bound proteins were denatured in Laemmli 2X for 5 min at 95°C and subjected to immunoblot analysis.

#### Split luciferase assay

The different T3Es-Nluc and Cluc-IDs fusion proteins were transiently expressed in *N. benthamiana* leaves using *Agrobacterium*-mediated transformation. After 48 hours, leaves were infiltrated with 1mM luciferin (XenoLight D-Luciferin, Perkin Elmer) and were imaged using a ChemiDoc imaging system (Bio-Rad). For quantification, single 4 mm punch disks were placed into wells of a 96 well plate, washed with sterile water and incubated with 1mM luciferin, as previously described [41, 43]. Briefly, after 10 min incubation, light emission was quantified in each well for 5 sec using a luminometer (PerkinElmer, VICTOR Nivo). For each interaction tested, 16 technical replicates were performed (16 leaf disks/plate) and this assay was replicated three times independently. As positive control, Cluc-MskA and RipG7-Nluc fusion proteins were used, as previously described [32].

#### Protein extraction and western blotting

*N. benthamiana* leaf discs were ground in liquid nitrogen and proteins were extracted in Laemmli buffer 2X. Yeast protein were extracted after a lysis step in 0.1 M NaOH during 20 min at room temperature followed by protein extraction in Laemmli buffer 1X. Immunodetection of proteins were performed by loading the samples on precast SDS-PAGE gels (4-15%, Bio-Rad). Proteins were transferred on nitrocellulose membrane using the Trans-Blot Turbo transfer system (Bio-Rad). The nitrocellulose membranes were blocked with 1X Tris-buffered Saline with Tween20 (TBS-T) solution (137 mM NaCl, 0.1% Tween-20, 3% Milk). Proteins transferred on nitrocellulose membranes were stained in Ponceau S staining solution (0.5% Ponceau S (w/v), 1% acetic acid). Immuno-detection of the protein of interest were performed with the following antibodies: anti-GFP (1:3000, mouse mAb (clones 7.1 and 13.1), Sigma), anti-HA-HRP (1:5000, Rat mAb (clone 3F10), Sigma), anti-Gal4-AD (1:5000, mouse mAb, Takara), Anti-Ac-K (1:2000, mouse mAb Ac-K-103, Cell signaling), anti-His6-HRP (1:50000, mouse mAb (clone His-2), Roche), anti-luciferase (1:10000, L0159, Sigma), anti-rabbit IgG-HRP (1:10000, Cell signaling) and polyclonal goat anti-mouse-Hrp (1:10000, Bio-Rad).

#### Arabidopsis transformation

The pDEST pB7FWG2-Δ35S (derived from pB7FWG2 from which 35S promoter sequence has been excised) was used for Arabidopsis transformation. Mutation of the WKRYGQK heptad of RRS1-R to the divergent WRKYGKR heptad was generated by site-directed mutagenesis using PrimeStar HS DNA polymerase from Takara Bio Inc. (Otsu, Japan). The following nucleotide substitutions were done: RRS1-R-Q1220K/K1221R (codon 1220: CAA to AAA and codon 1221:AAA to AGG). pENTR221 clones containing either [RPS4 and RRS1-R^Ws-2^] or [RPS4 and RRS1-R^WRKYGKR^] genes under their genomic 5’ and 3’ regulatory sequences were used for LR recombination with pDEST pB7FWG2-Δ35S plasmid. The *rps4-21 rrs1-1* mutant [44] was transformed as previously described [45]. Transgenic T1 plants were selected on MS-media in presence of 10 µg/mL of Phosphinotricin (Duchefa Biochemie).

#### Plant inoculation with *Ralstonia solanacearum*

*A. thaliana* plants were inoculated by soil-drenching with a *R. solanacearum* (strain GMI1000) bacterial suspension at 5.10^7^ cfu/ml, as described previously [46]. The plants were then incubated in a growth chamber at 16h light at 27°C and 8 hours dark at 26°C. Disease development was recorded daily using a macroscopic scale describing the observed wilting: 1 for 25% of the leaves wilted; 2 for 50%; 3 for 75% and 4 for complete wilting. For subsequent analysis the data was transformed into a binary index: 0 for < 50% of wilted leaves and 1 for more or equal to 50% wilted leaves. To compare the disease development of two given strains, we used the Kaplan– Meier survival analysis with the Gehan–Breslow–Wilcoxon method to compute the P-value to test the null hypothesis of identical survival experience of the two tested strains [47]. A P-value smaller than 0.05 was considered significant. Statistical analyses were done with Prism version 9 (GraphPad Software). Hazard ratio calculation and representation were also done using Prism version 9 as described previously [32].

## ACCESSION NUMBERS

The accession numbers for the NLRs from which the IDs where cloned are: At5G17890.1, At5G45050.2, At4G12020.3, At5G45260.1, Glyma.05G165800.4.p, At5G66630.1, At5G17890.1, At4G16990.2, At4G16990.1, At4G12020.3, At4G12020.1, MDP0000457940, Bradi4g24914, At4G19500.2, At5G47260.1, Solyc04g008200.2.1, Solyc05g005130.1.1, Solyc04g007490.2.1, Solyc05g012890.1.1, Solyc10g008220.2.1, Solyc05g013260.1.1, Solyc06g065000.1.1, Solyc05g007170.2.1, Solyc05g007640.2.1, Solyc05g008650.1.1, Solyc05g012910.2.1, Solyc04g008180.1.1, Solyc05g012740.1.1, Solyc05g005330.2.1, Solyc05g007350.1.1, Solyc10g051170.1.1, Solyc09g072940.1.1, Solyc05g012740.1.1, Solyc01g102880.1.1, Medtr3g030980.1, Medtr3g032760.1, Medtr3g015260.2, Medtr3g018930.2, Medtr6g087200.1, Medtr6g015490.1, Medtr6g015665.1, MDP0000286727, MDP0000457940, Bradi3g34961.1.p, Bradi3g58937.5.p, LOC_Os02g19890.3, LOC_Os11g11920.1, LOC_Os07g17230.1, LOC_Os05g15040.1, LOC_Os12g18360.1, LOC_Os08g30634.1, LOC_Os09g20020.1, LOC_Os11g11810.1, LOC_Os11g46210.1, Glyma.12G236500.1.p, Glyma.12G236500.1.p. This list is also detailed in Table 1.

## ACKNOWLEDGEMENTS

We thank Thomas Kroj for providing *O. sativa* cv nipponbare cDNA, Armin Djamei for *B. distachyon* Bd21, *G. max* Wm82 cDNA, Arry Morel for *S. lycopersicum* M82 cDNA, Marie-Françoise Jardinaud for *M. truncatula* A17 cDNA and Emilie Vergne for *M. domestica* cDNA. This work was supported by the French Laboratory of Excellence project ‘‘TULIP’’ (ANR-10-LABX-41; ANR-11-IDEX-0002-02). No conflict of interest are known to the authors at the moment of submitting this work.

## SHORT LEGENDS FOR SUPPORTING INFORMATIONS

**Figure S1**. Integrated domains (IDs) cloning strategy

**Figure S2**. Number of ID identified, selected and successfully cloned

**Figure S3**. Yeast cells co-expressing ID#29 with several *R. solanacearum* T3Es are able to grow on selective media

**Figure S4**. Immuno-detection of the different Gal4-(AD)-IDs fusion proteins in yeast

**Figure S5**. Immuno-detection of T3E-Nluc and Cluc-ID fusion proteins transiently expressed in *N. benthamiana* cells.

**Figure S6**. GST pull-down assay showing that WRKY-ID#85-3HA forms a complex with GST-PopP2-6His.

**Figure S7**. Schematic representation of the different domains present in RRS1-R and GmNLR-ID#85.

**Figure S8**. Bacterial and *in planta* acetylation assays show that WRKY-ID#85 does not behave as a substrate of PopP2 acetyltransferase.

**Figure S9**. Introduction of the WRKY-ID#85 divergent heptad in RRS1-R receptor results in loss of responsiveness to PopP2.

**Figure S10**. NLR-ID#85 orthologs are prone to harbor WRKY as atypical domain.

**Figure S11.** Original raw Y2H matrices. All yeast growth spots used in Figure 1 and Figure S3 and circled by white dots. The Red circle represents an positive control. RipG7* (notedG7*) is a plant recoded version of RipG7 (noted G7). All Integrated domains and *R. solanacearum* effectors are indicated in the matrices.

## TABLES

**Table S1:**
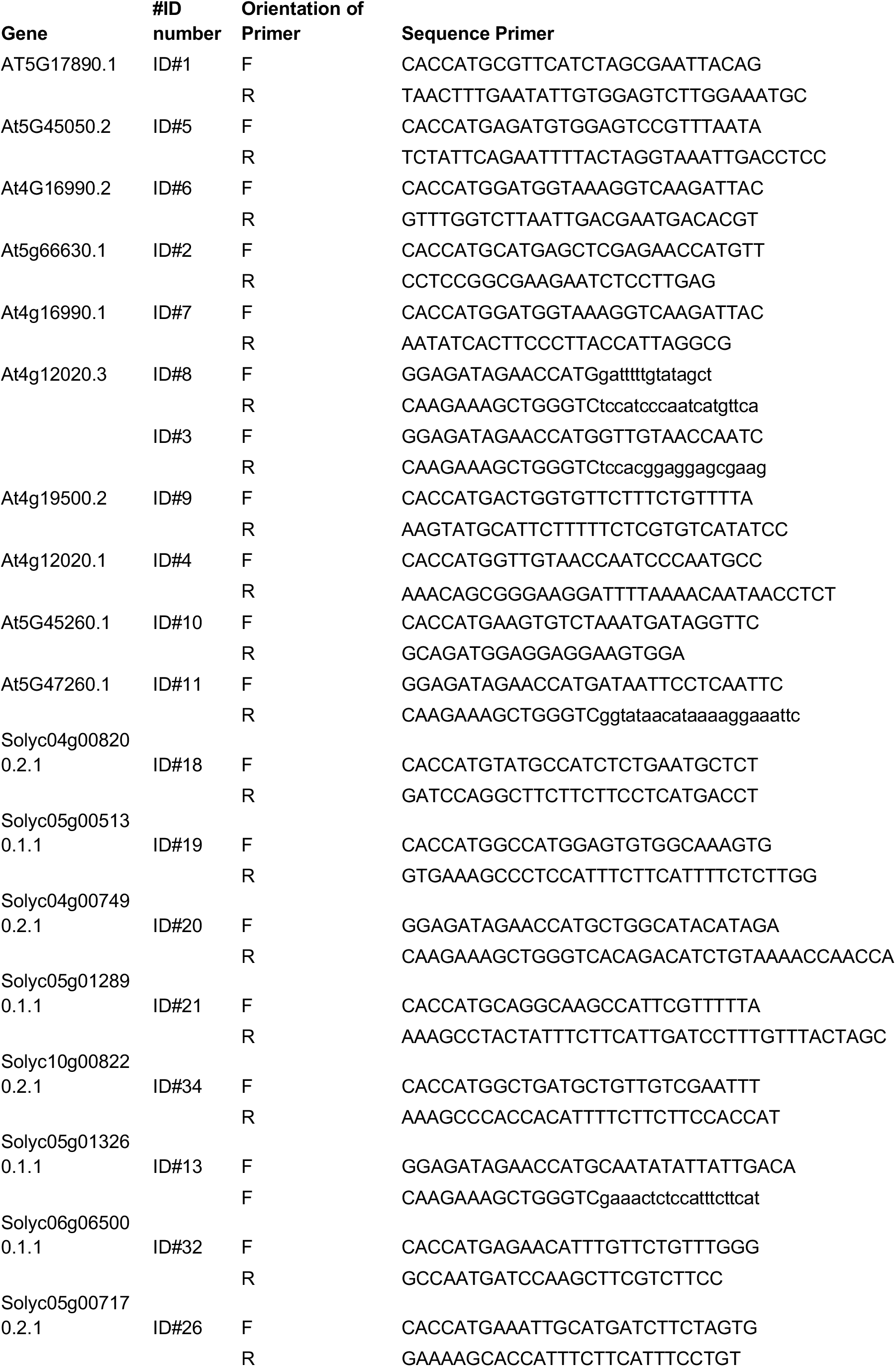
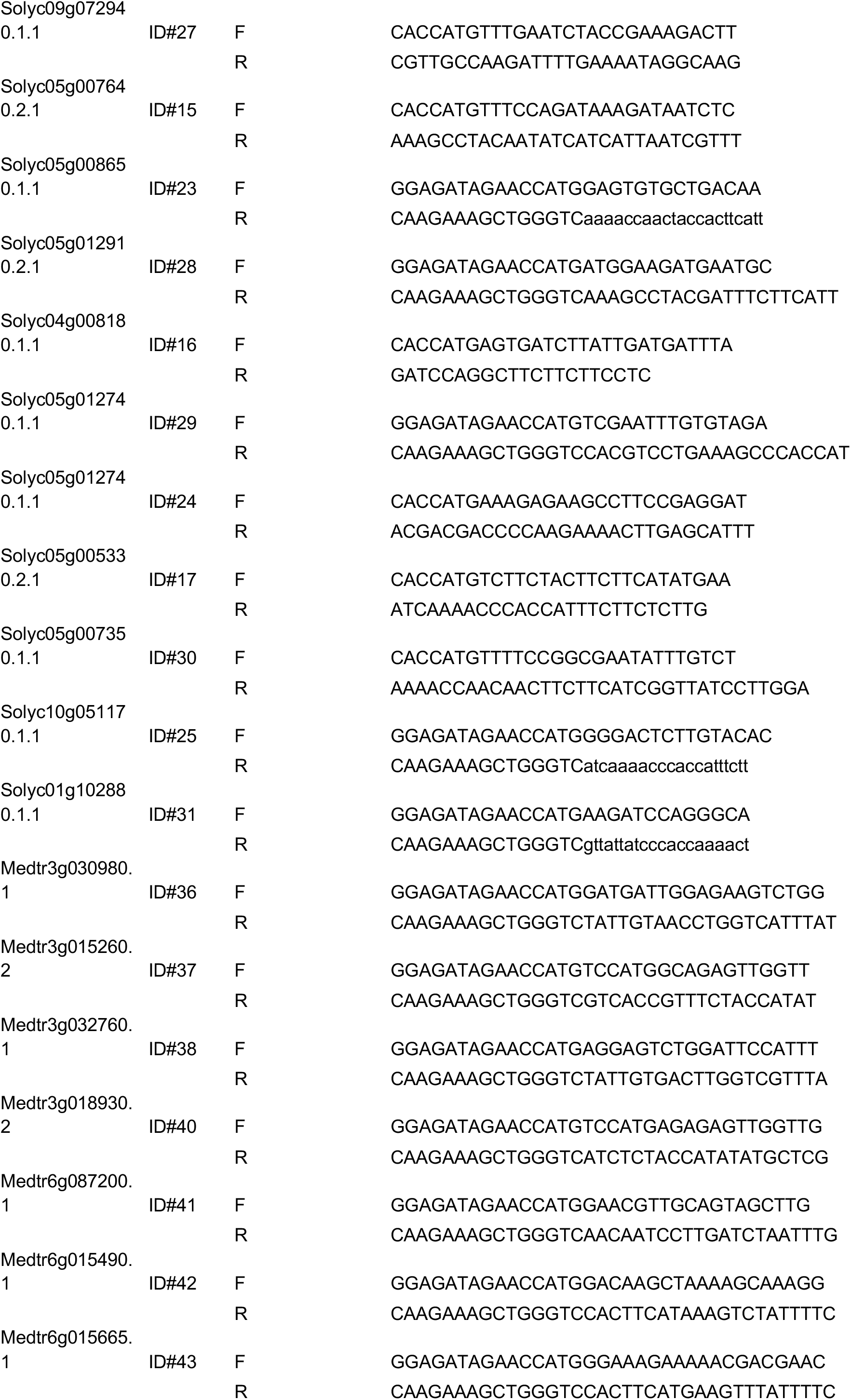
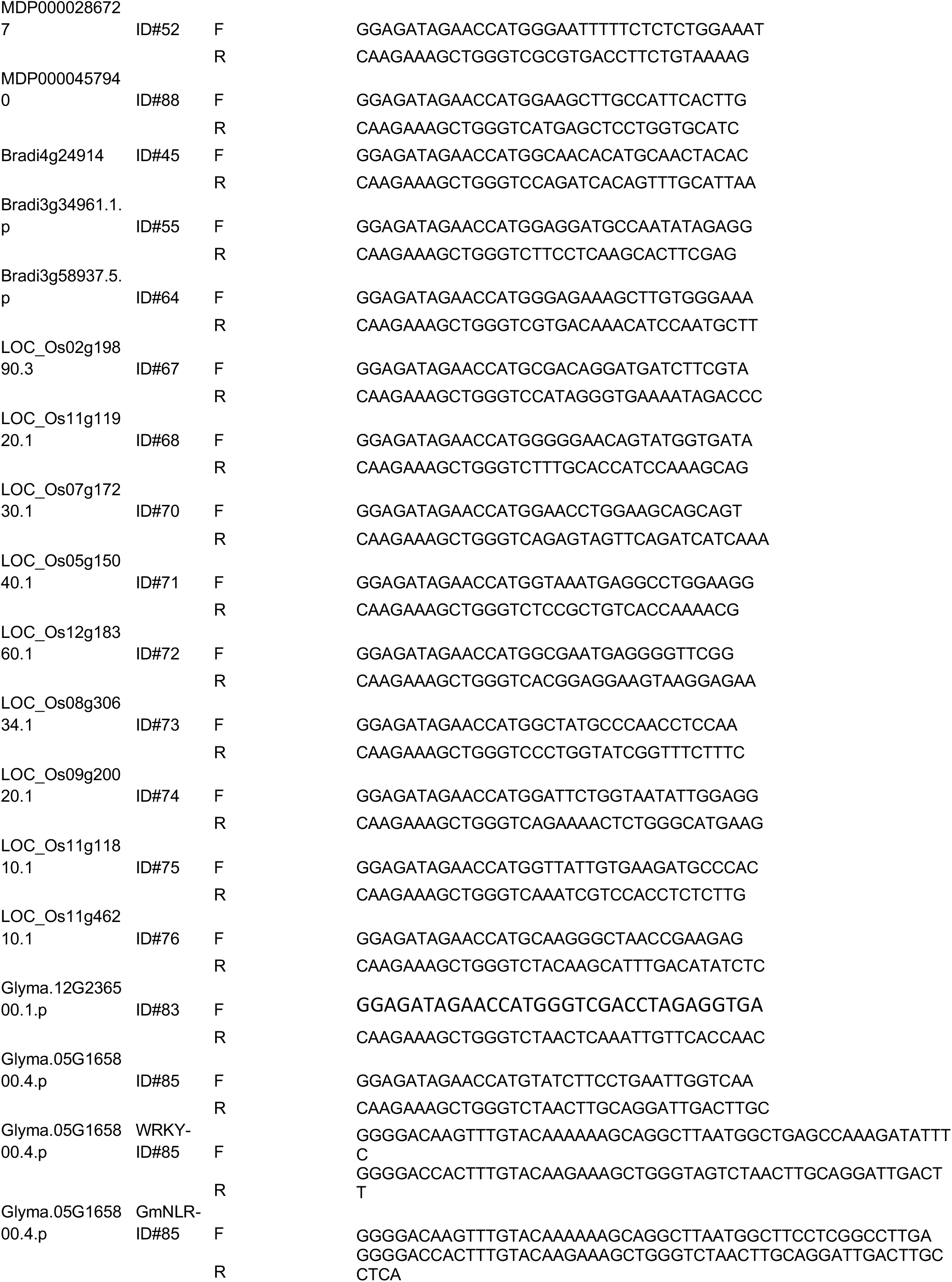
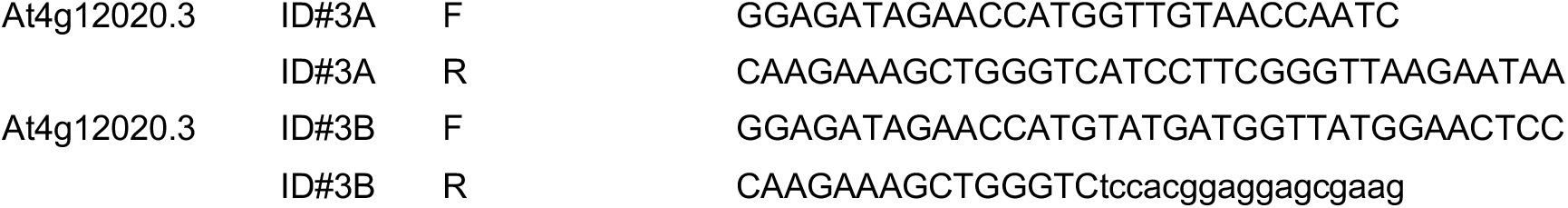
List of primers used in this study.

