## Supplementary Figures and tables for "An NLR Integrated Domain toolkit to identify plant pathogen effector targets"

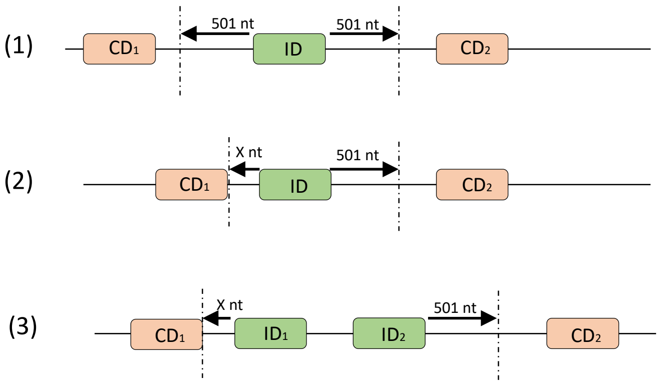


**Figure S1. Integrated Domains (IDs) cloning strategy**

Each ID clone was generated by following a specific strategy based on three possibilities : (1) An isolated ID is cloned with 501 upstream and 501 downstream flanking nucleotides. (2) the ID surrounded by canonical domains is cloned at the border of these domains, X being a multiple of 3 to maintain the in-frame fusion. (3) two IDs following each other are cloned together.

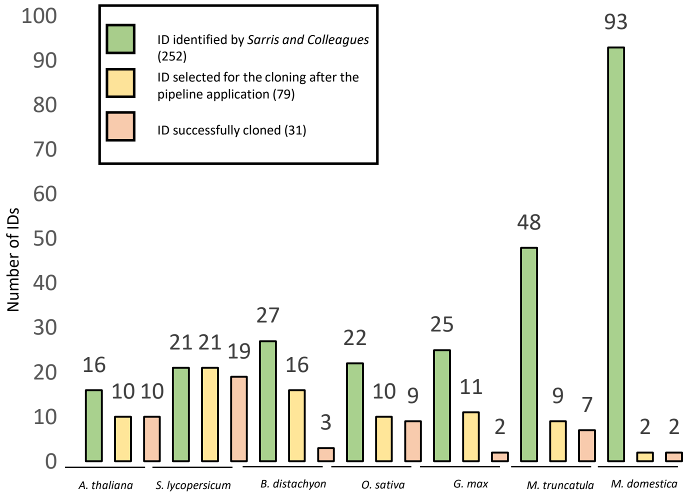


**Figure S2. Number of IDs identified, selected and successfully cloned.** The histograms show the reduction of the number of IDs after the application of our stringent pipeline (green versus yellow histograms). Orange histograms represent the number of IDs successfully cloned. On the 252 IDs identified by Sarris and colleagues, 79 IDs were selected and 52 of them were cloned, representing 31 Pfam domains (see Table 1). For *Medicago truncatula* (*M. truncatula*) and Malus domestica (*M. domestica*), miss-annotation of the *NLR-ID* gene models strongly reduced the number of IDs selected for cloning.

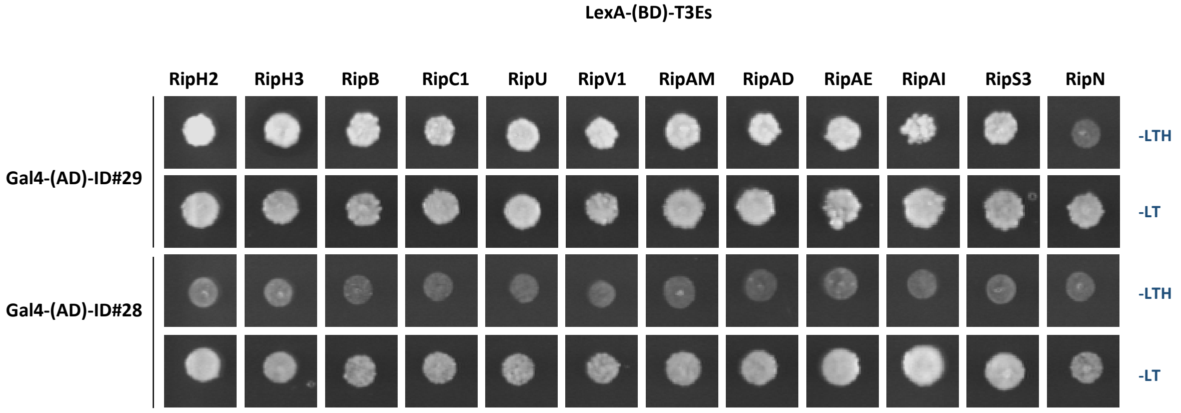


**Figure S3: Yeast cells co-expressing ID#29 with several *R. solanacearum* T3Es are able to grow on selective media.**
Co-expression of the Gal4-(AD)-ID#29 fusion protein with RipH2, RipH3, RipB, RipC1, RipU, RipV1, RipAM, RipAD, RipAE, RipAI, and RipS3 effectors fused with LexA-(BD) domain confers prototrophy of yeast cells for growth on selective media (-LTH). In this assay, LexA-(BD)-RipN and Gal4-ID#28 were used as negative bait and prey controls, respectively. The screening was conducted two times with similar results (two technical replicates) and the different yeast colonies were plated on SD–LT selective media for selection of mated cells.

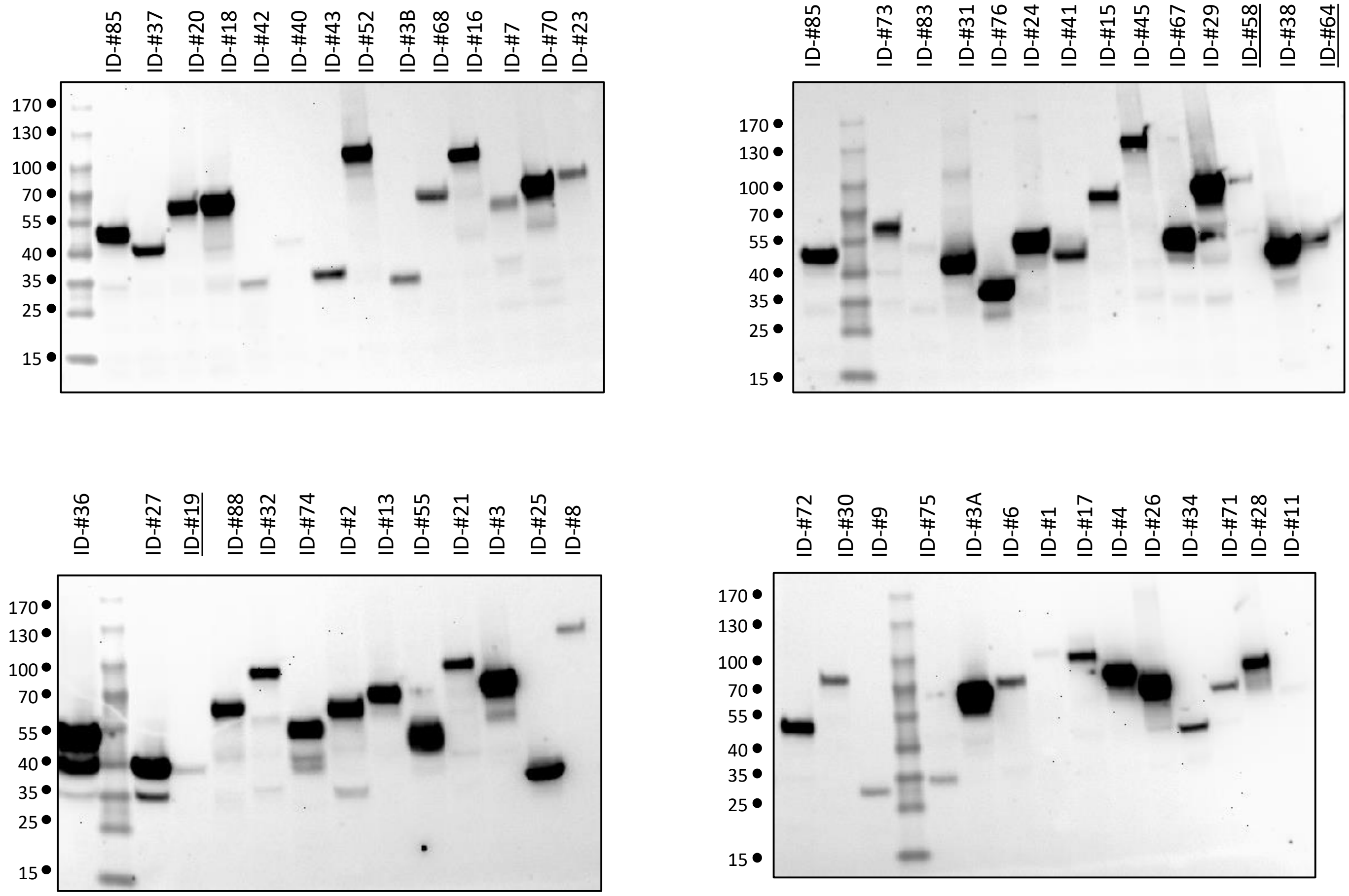


**Figure S4. Immuno-detection of the different Gal4-(AD)-IDs fusion proteins in yeast**

Total yeast proteins were extracted from diploïd yeast cells and subjected to immunoblot analysis. The different Gal4-(AD)-IDs fusion proteins were detected using an antibody raised against the Gal4-(AD) domain (primary antibody). Some ID# could not be detected in yeast with correct expected size (underlined: ID#58, #64, #19).

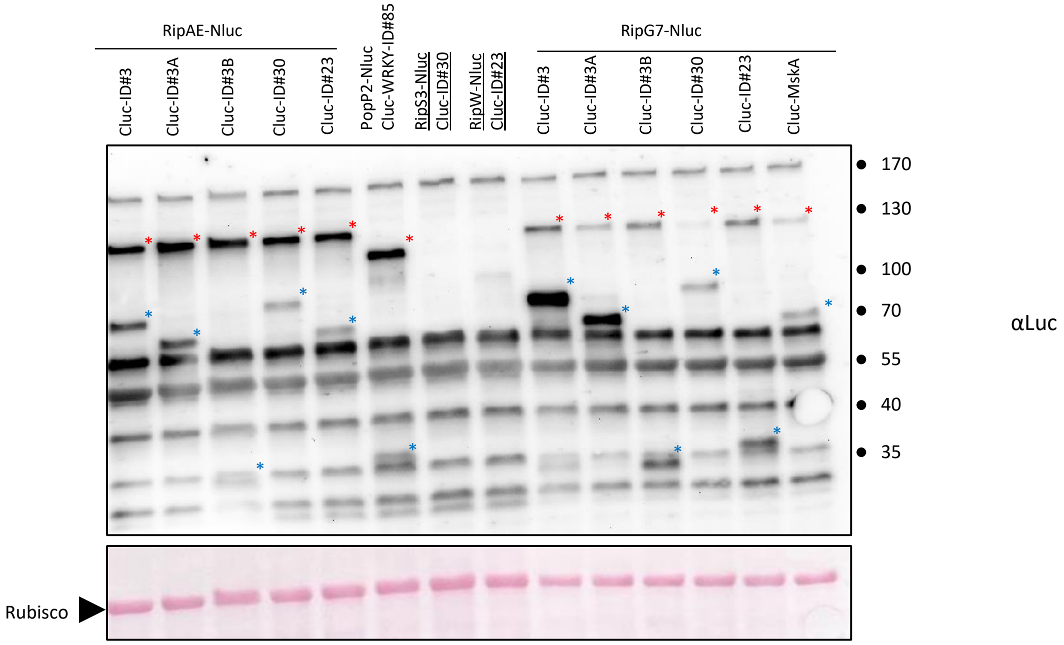


**Figure S5. Immuno-detection of T3E-Nluc and Cluc-ID fusion proteins transiently expressed in *N. benthamiana* cells.**
Expression of T3E-Nluc and Cluc-ID proteins *in planta* was checked by immunoblot with an anti-luciferase antibody. For all co-expressed proteins, signals corresponding to the size of both fusion proteins were detected with the exception of RipS3-Nluc co-expressed with Cluc-ID#30 and RipW-Nluc co-expressd Cluc#23 for which no signal was detected (underlined). Although Cluc-ID#23 and Cluc-ID#30 proteins were immuno-detected in other co-expressions. Red stars and blue stars indicate Nluc- and Cluc- fusion proteins, respectively.

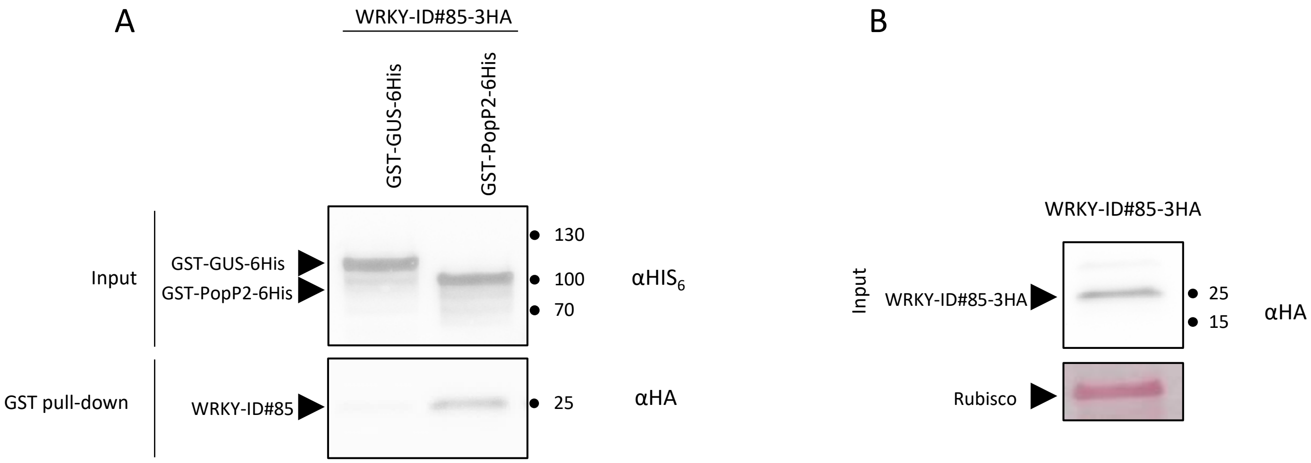


**Figure S6. GST pull-down assay showing that WRKY-ID#85-3HA forms a complex with GST-PopP2-6His.**
(A) A GST pull-down assay was peformed by incubating recombinant GST-GUS-6His or GST-PopP2-6His proteins bound on Glutathione beads with a plant protein extract from *N. benthamiana* cells transiently expressing WRKY-ID#85-3HA (see B). GST- and HA-tagged proteins were immunodetected with anti-His6 and anti-HA antibodies, respectively. WRKY-ID#85-3HA was able to form a complex with GST-PopP2-6His bound on Gluthatione beads, but not with GST-GUS-6His. (B) Immuno-detection of WRKY-ID#85-3HA expressed in *N. benthamiana* cells with an anti-HA antibody. An equal volume of this *N. benthamiana* protein extract was incubated with recombinant GST-GUS-6His or GST-PopP2-6His proteins bound on Gluthathione beads.

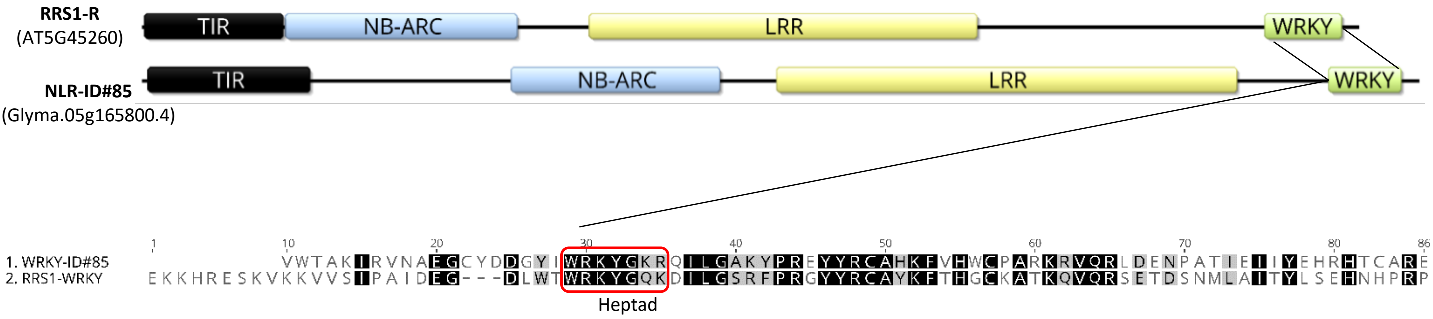


**Figure S7: Schematic representation of the different domains present in RRS1-R and GmNLR-ID#85.** Protein domains were identified by Interpro scan. Comparison of RRS1-R and GmNLR-ID#85 structure shows that their WRKY domain is integrated in the C-terminal portion of the proteins. The divergent WRKY heptad in GmNLR-ID#85 is highlighted (rex box) in the alignement between the WRKY domains of RRS1-R (residues 1190 to 1272) and GmNLR-ID#85 (residues 1258 to 1334). The protein alignment was performed with Blosum62 matrix (threshold 1) by Geneious software (Geneious 11.1.4).

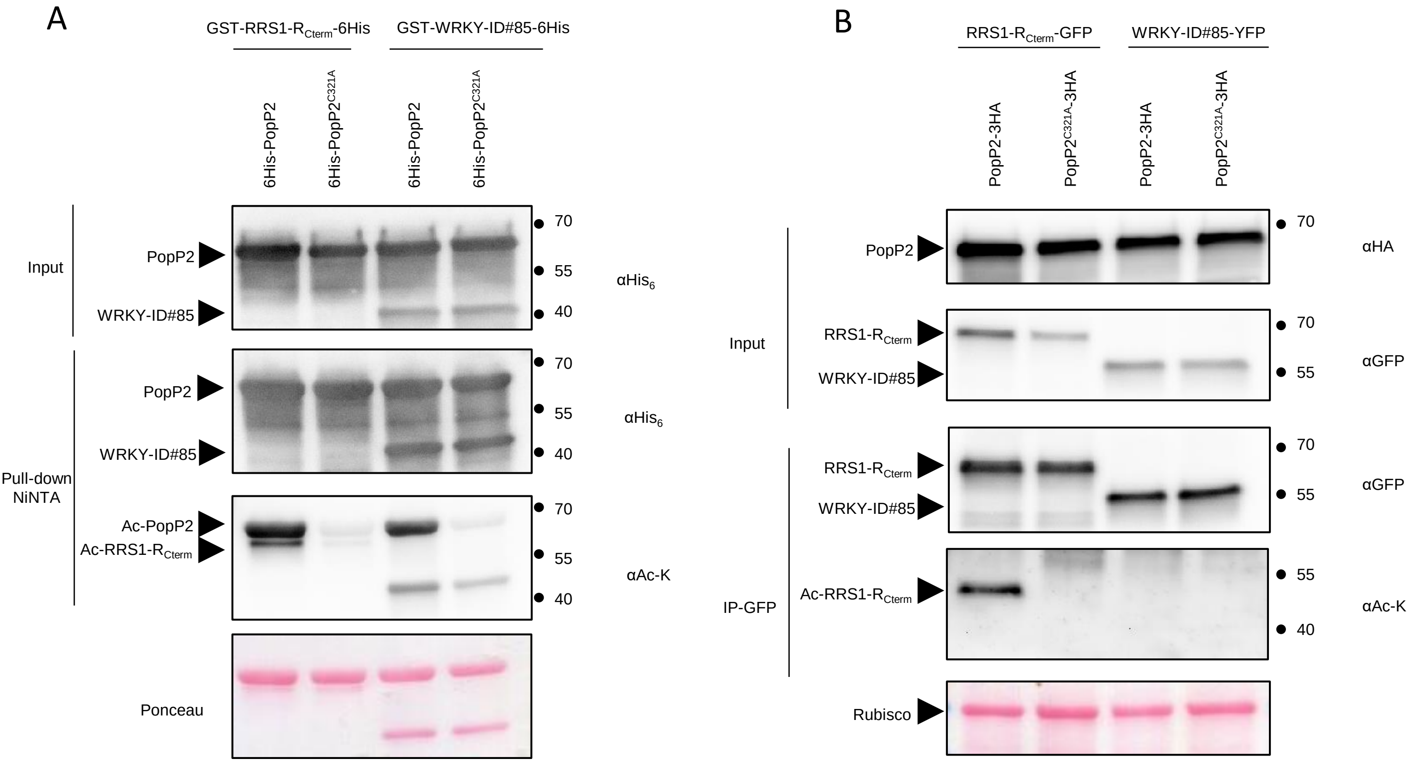


**Figure S8: Bacterial and *in planta* and acetylation assays show that WRKY-ID#85 does not behave as a substrate of PopP2 acetyltransferase.** (A) GST-RRS1-$R_{Cterm}$-6his and GST-WRKY-ID#85-6His were co-expressed in *E. coli* either with wild-type PopP2 or its catalytic mutant, both of them N-terminally fused with a 6His tag. All 6His-tagged proteins were bound on Ni-NTA resin. Lys-acetylated proteins were immuno-detected with an anti-acetyl lysine (a-Ac-K) antibody. Immunoblots shows that, unlike GST-RRS1-$R_{Cterm}$-6His, GST-WRKY-ID#85-6His co-expressed with active 6His-PopP2 is not recognized by an anti-Ac-K antibody, suugesting that WRKY-ID#85 does not behave as a substrate of PopP2 enzymatic activity.

(B) RRS1-$R_{Cterm}$-eGFP and WRKY-ID#85-YFP were transiently expressed in *N. benthamiana* either with active PopP2-3HA or catalytically inactive PopP2$^{C321A}$-3HA. Both fluorescent fusion proteins were immunoprecipitated on GFP beads and subjected to immunoblot analysis using anti-GFP and anti-Ac-K antibodies. As shown in Fig S8.A, RRS1-$R_{cterm}$ but not WRKY-ID#85GFP is acetylated by active PopP2. Ponceau S staining indicates that similar amounts of protein were loaded in the different lanes.

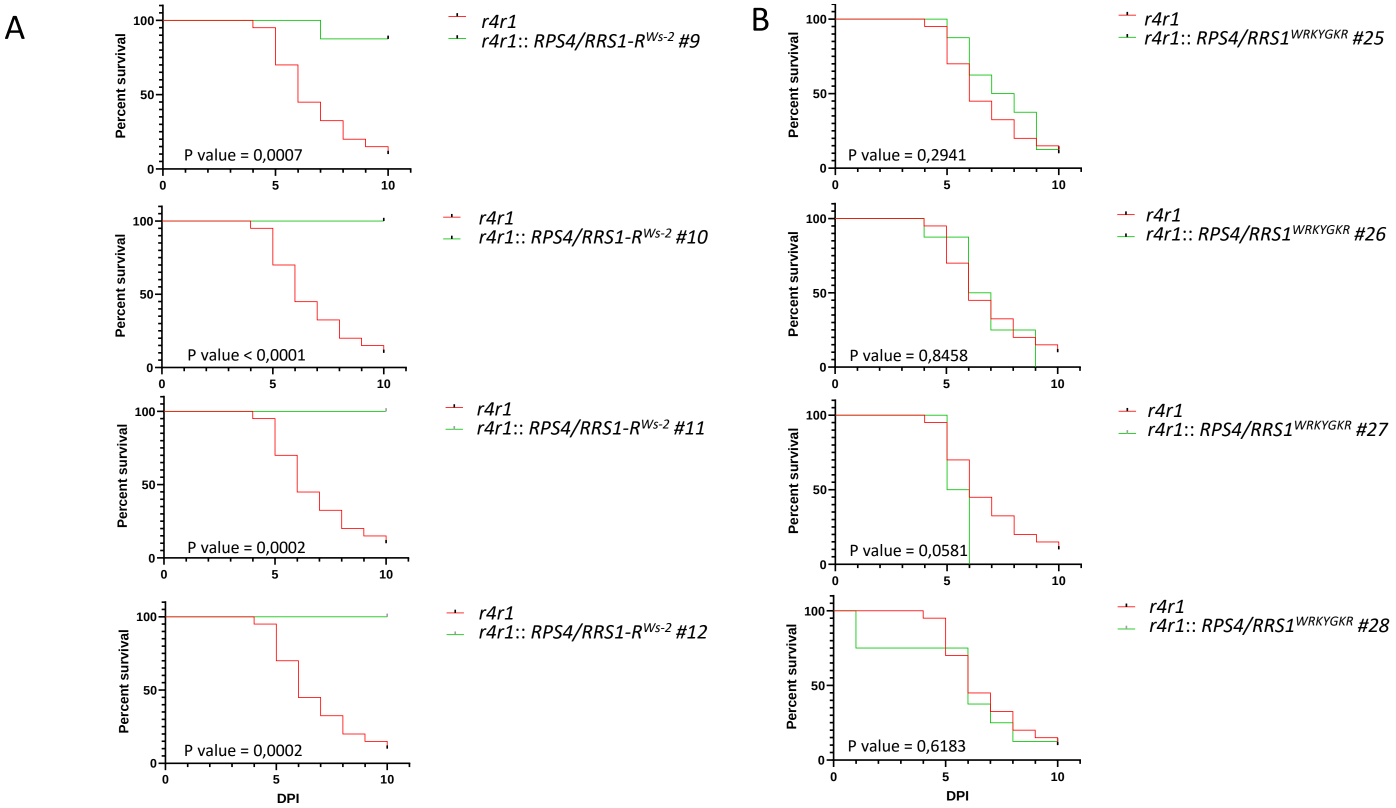


**Figure S9. Introduction of the WRKY-ID#85 divergent heptad in RRS1-R receptor results in loss of responsiveness to PopP2.**
(A) Four T2 transgenic lines (#9 to #12) expressing RPS4 and RRS1-R genes from Ws-2 under their genomic 5' and 3' regulatory sequences were made in *rps4-21 rrs1-1* double mutant (Ws-2 background, hereinafter designated by *r4r1*). These lines were root inoculated with *R. solanacearum* GMI1000 strain. All the tested lines displayed a resistance phenotype at 10 dpi. As indicated in the survival analysis and their associated logrank test pValue allowing to reject the H0 hypothesis of curve similarity.
(B) The divergent heptad WRKYGKR present in WRKY-ID#85 was introduced in the WRKY domain of RRS1-R (RRS1-R$^{WRKYGKR}$). Four T2 lines (#25 to #28) expressing RPS4 and RRS1-R$^{WRKYGKR}$ isoforms under their genomic 5' and 3' regulatory sequences were made in *rps4-21 rrs1-1* double mutant and were root-inoculated with *R. solanacearum* GMI1000 strain. All the tested lines were susceptible to GMI1000 and wilted like the *r4r1* control line (unable to reject similarity of curve H0 hypothesis). Gehan-Breslow-Wilcoxon was used as statistical test in the graphpad prsim9 software.

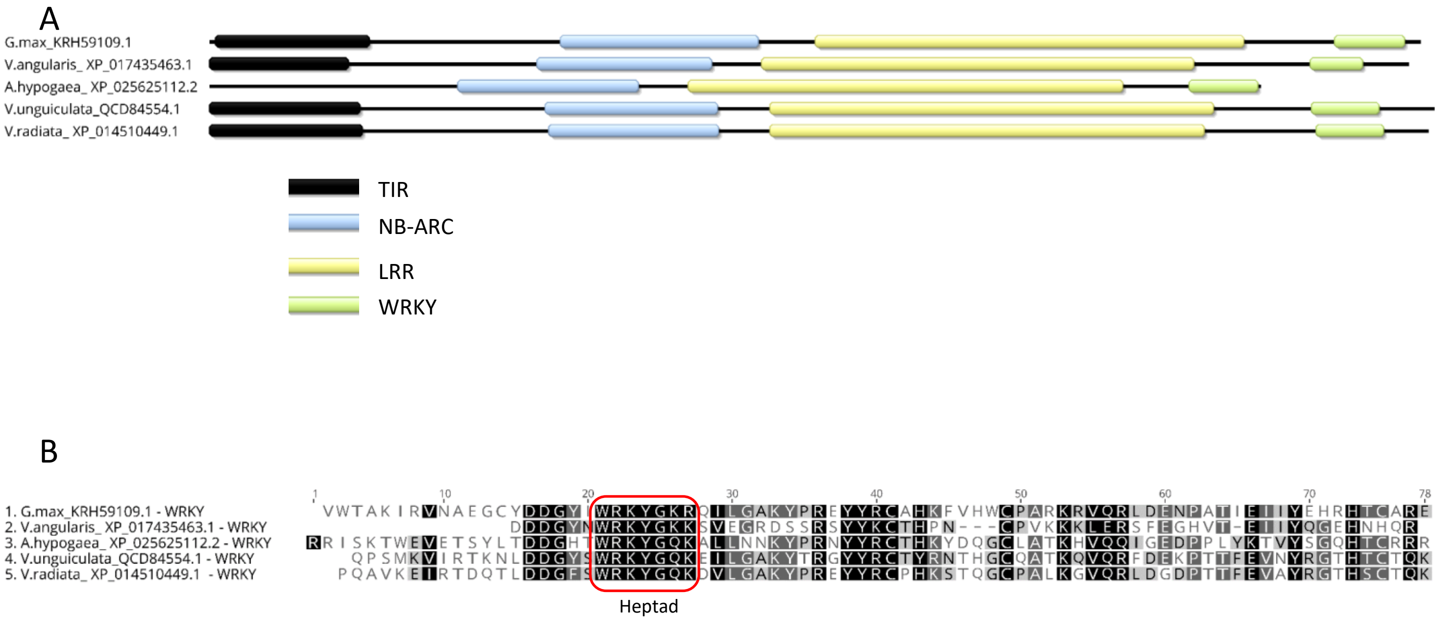


**Figure S10. NLR-ID#85 orthologs are prone to harbor WRKY as atypical domain.**
The GmNLR-ID#85 (G.max_KRH59109.1) sequence was used as bait to identify by blast orthologs in Fabaceae plant family. (A) Four NLRs have been identified from the following plant species: *Vigna angularis*, *Arachis hypogaea*, *Vigna unguiculata* and *Vigna radiata*. The interpro annotation allow us to identify conserved protein domains and WRKY domains as atypical domains. (B) The WRKY domains alignment reveals that *Glycine max* and *Vigna angularis* proteins carry a mutated WRKY heptad (respectively, WRKYGKR and WRKYGKK) while the others have the well conserved WRKY heptad (WRKYGQK). The protein alignment was performed with Blosum62 matrix (threshold 1) by Geneious software (Geneious 11.1.4).

**Figure S11**. Original raw Y2H matrices. All yeast growth spots used in Figure 1 and Figure S3 and circled by white dots. The Red circle represents an positive control. RipG7* (notedG7*) is a plant recoded version of RipG7 (noted G7). All Integrated domains and *R. solanacearum* effectors are indicated in the matrices.

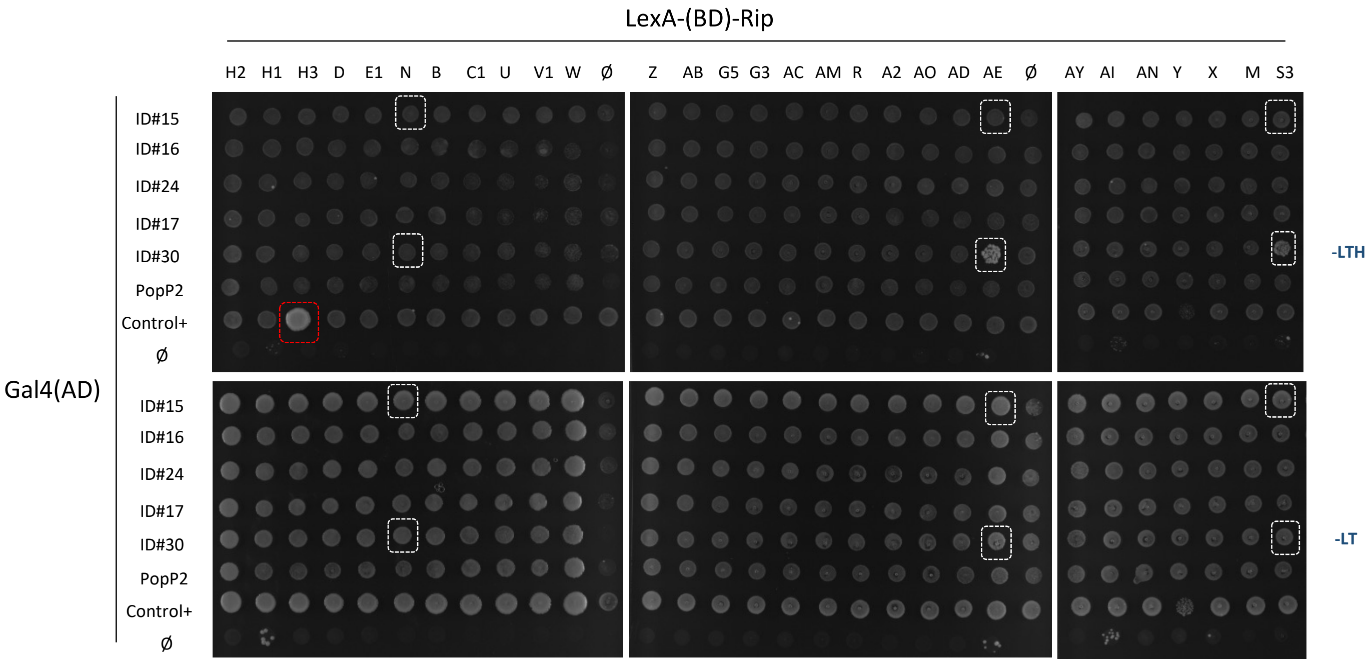

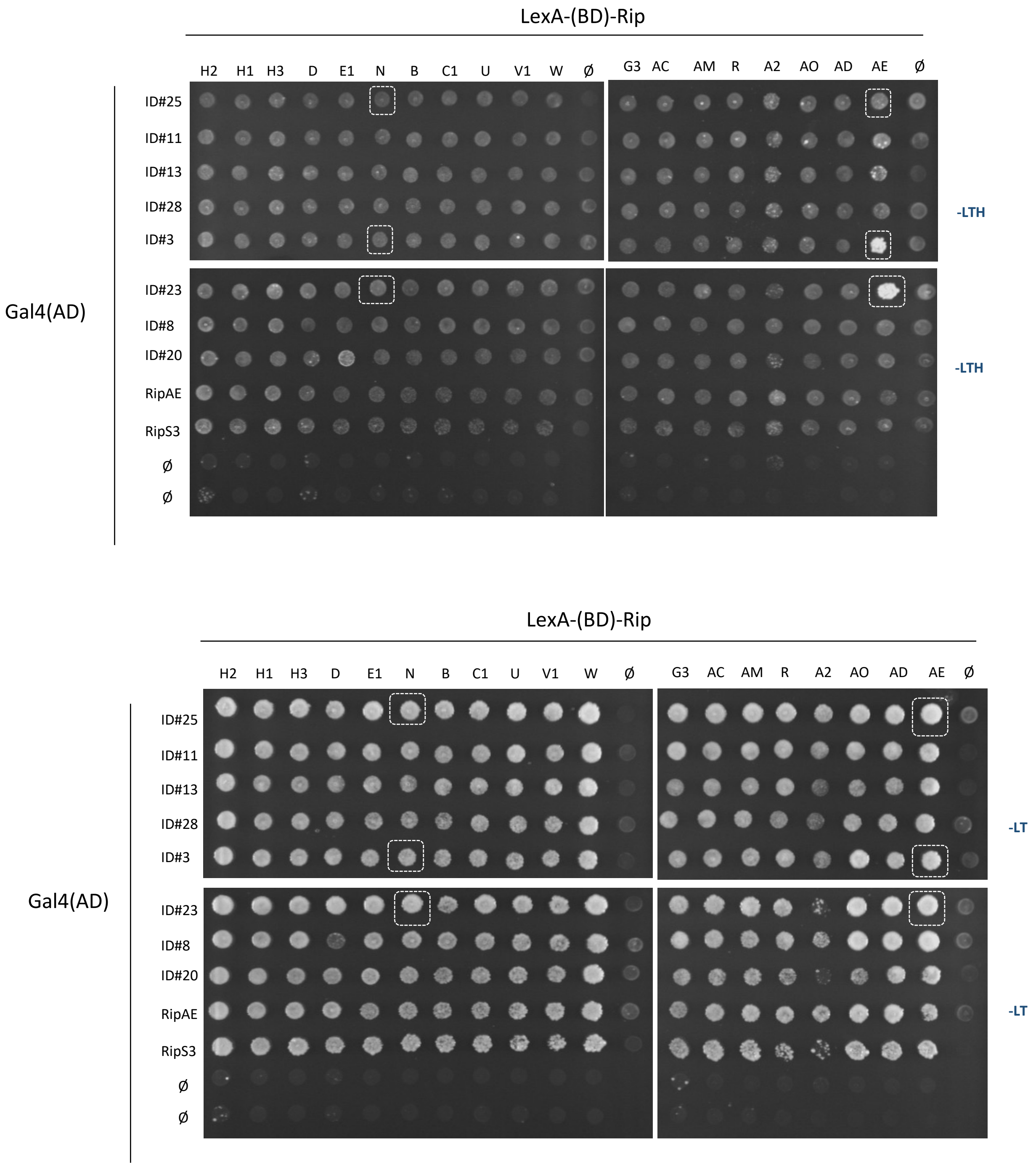
LexA-(BD)-Rip
H2 H1 H3 D E1 N B C1 U V1 W Ø G3 AC AM R A2 AO AD AE Ø
Gal4(AD)
ID#25
ID#11
ID#13
ID#28
ID#3
ID#23
ID#8
ID#20
RipAE
RipS3
Ø
Ø
-LTH
-LTH
LexA-(BD)-Rip
H2 H1 H3 D E1 N B C1 U V1 W Ø G3 AC AM R A2 AO AD AE Ø
Gal4(AD)
ID#25
ID#11
ID#13
ID#28
ID#3
ID#23
ID#8
ID#20
RipAE
RipS3
Ø
Ø
-LT
-LT

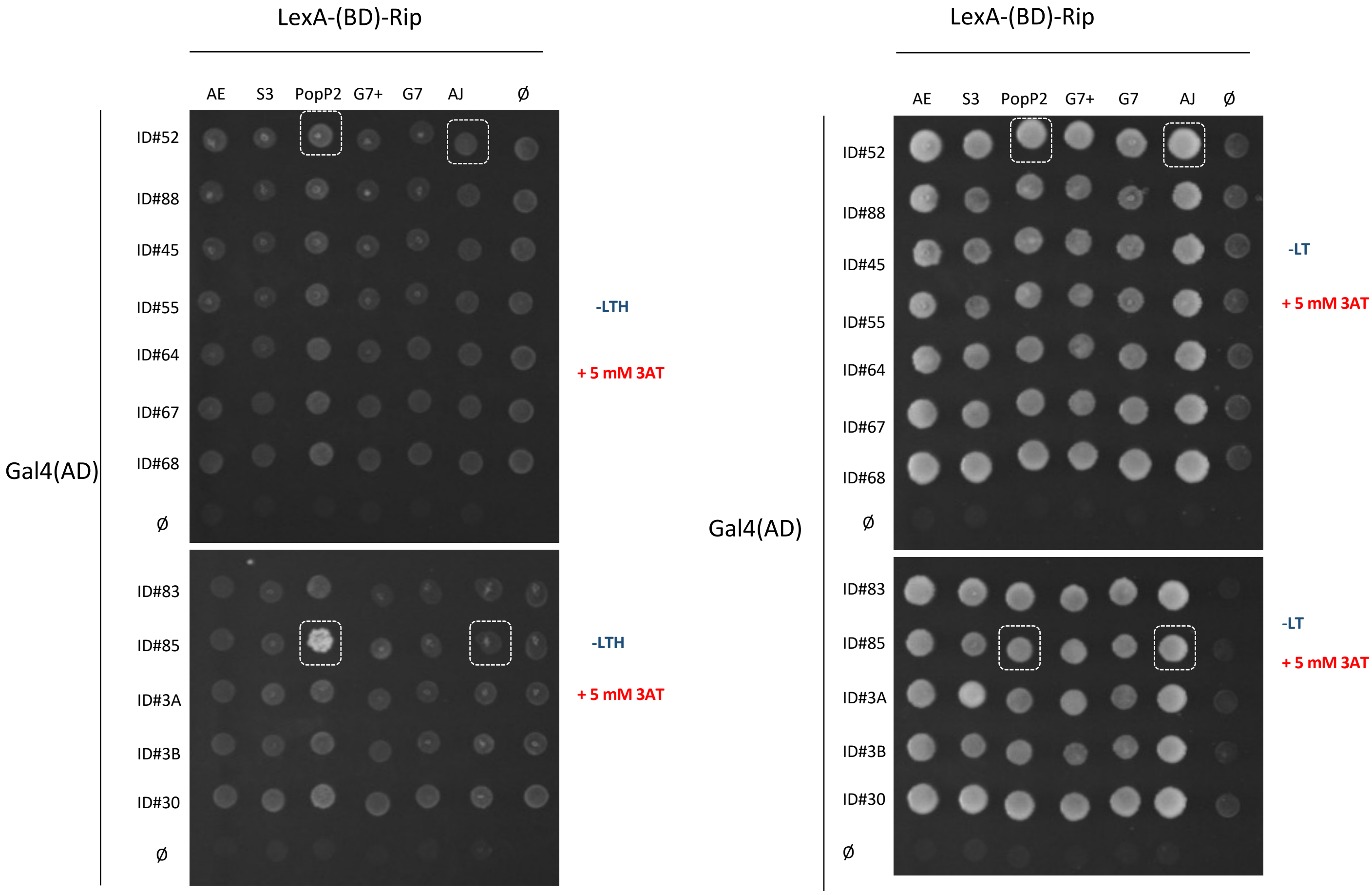

LexA-(BD)-Rip
AE S3 PopP2 G7+ G7 AJ Ø
Gal4(AD)
ID#52
ID#88
ID#45
ID#55
ID#64
ID#67
ID#68
Ø
ID#83
ID#85
ID#3A
ID#3B
ID#30
Ø
-LTH
+ 5 mM 3AT
-LTH
+ 5 mM 3AT
LexA-(BD)-Rip
AE S3 PopP2 G7+ G7 AJ Ø
Gal4(AD)
-LT
+ 5 mM 3AT
-LT
+ 5 mM 3AT

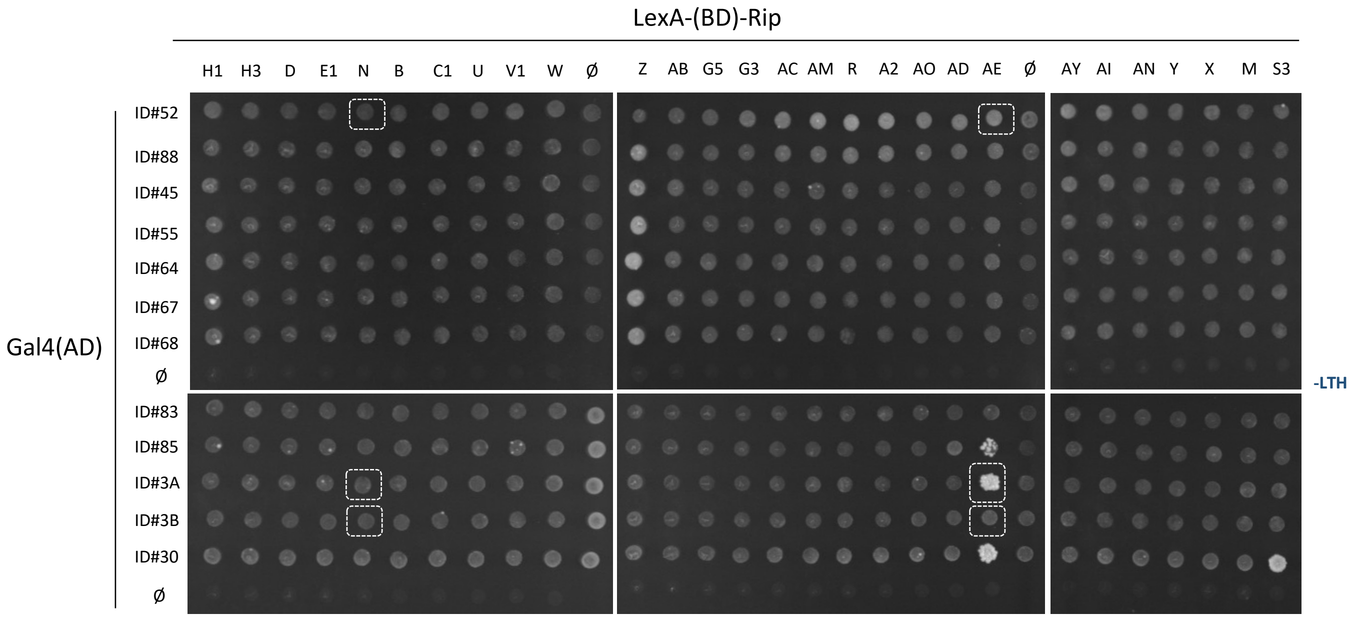
LexA-(BD)-Rip
H1 H3 D E1 N B C1 U V1 W Ø
Z AB G5 G3 AC AM R A2 AO AD AE Ø
AY AI AN Y X M S3
Gal4(AD)
ID#52
ID#88
ID#45
ID#55
ID#64
ID#67
ID#68
Ø
ID#83
ID#85
ID#3A
ID#3B
ID#30
Ø
-LTH

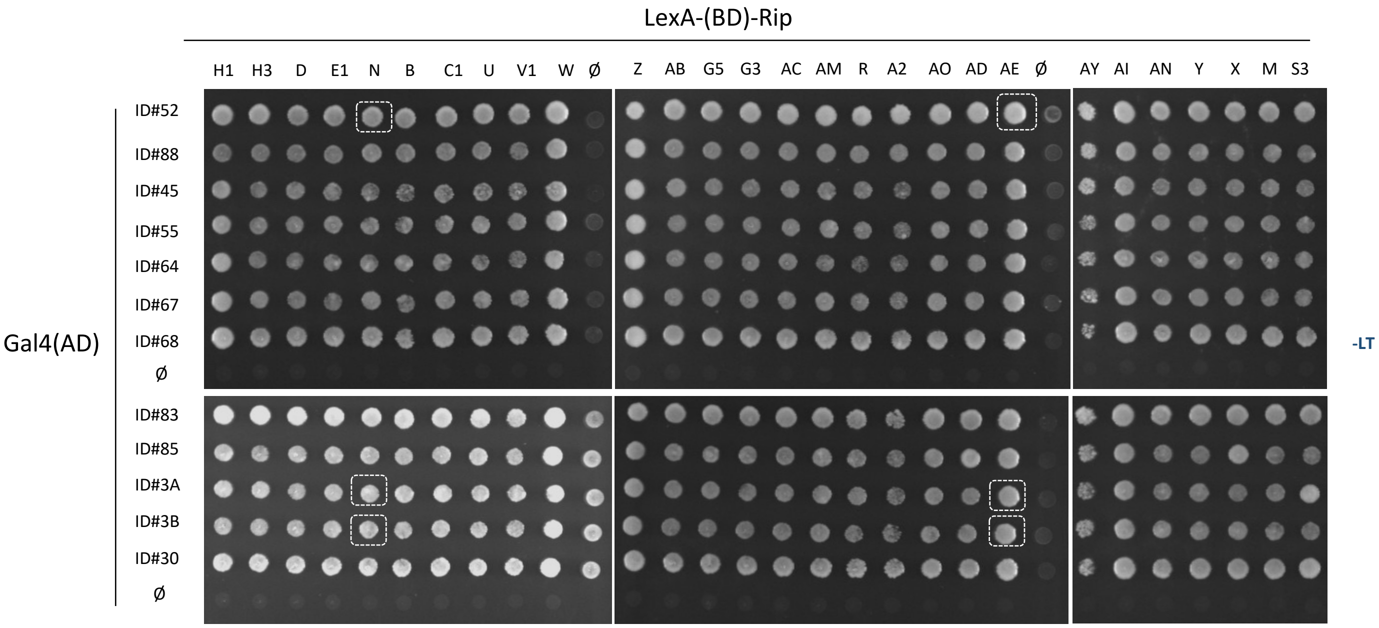
LexA-(BD)-Rip
H1 H3 D E1 N B C1 U V1 W Ø
Z AB G5 G3 AC AM R A2 AO AD AE Ø
AY AI AN Y X M S3
Gal4(AD)
ID#52
ID#88
ID#45
ID#55
ID#64
ID#67
ID#68
Ø
ID#83
ID#85
ID#3A
ID#3B
ID#30
Ø
-LT

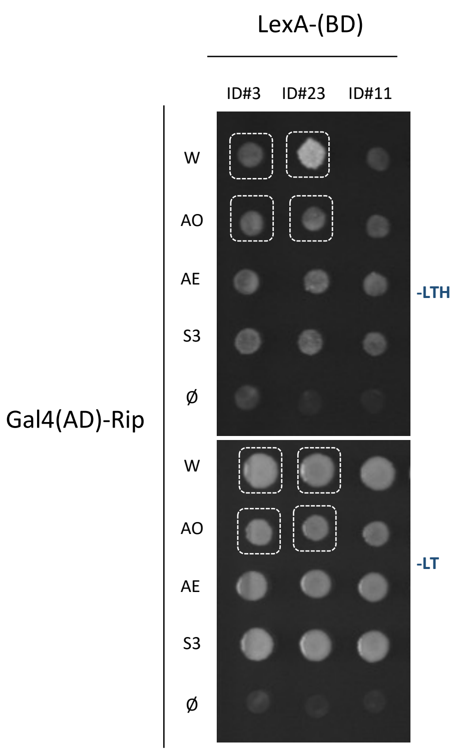

LexA-(BD)
ID#3 ID#23 ID#11
Gal4(AD)-Rip
W
AO
AE
S3
Ø
-LTH
W
AO
AE
S3
Ø
-LT


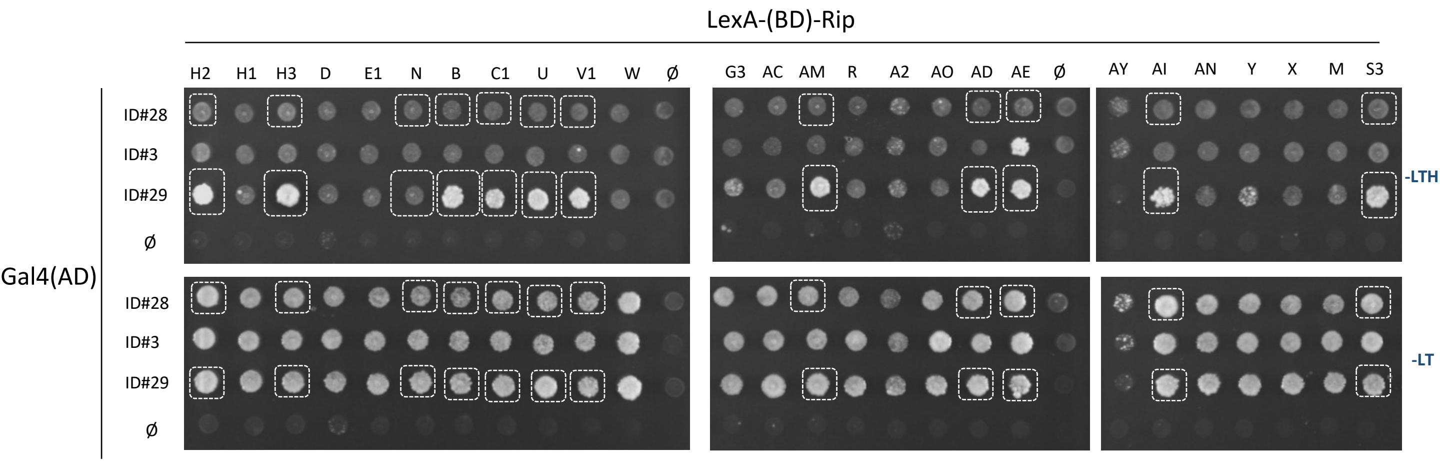

LexA-(BD)-Rip
H2 H1 H3 D E1 N B C1 U V1 W Ø
G3 AC AM R A2 AO AD AE Ø
AY AI AN Y X M S3
Gal4(AD)
ID#28
ID#3
ID#29
Ø
-LTH
ID#28
ID#3
ID#29
Ø
-LT

**Table S1: List of primers used in this study**

| Gene | #ID number | Primer orientation | primer sequence |
|---|---|---|---|
| AT5G17890.1 | ID#1 | F | CACCATGCGTTCATCTAGCGAATTACAG |
| | | R | TAACTTTGAATATTGTGGAGTCTTGGAAATGC |
| At5G45050.2 | ID#5 | F | CACCATGAGATGTGGAGTCCGTTTAATA |
| | | R | TCTATTCAGAATTTTACTAGGTAAATTGACCTCC |
| At4G16990.2 | ID#6 | F | CACCATGGATGGTAAAGGTCAAGATTAC |
| | | R | GTTTGGTCTTAATTGACGAATGACACGT |
| At5g66630.1 | ID#2 | F | CACCATGCATGAGCTCGAGAACCATGTT |
| | | R | CCTCCGGCGAAGAATCTCCTTGAG |
| At4g16990.1 | ID#7 | F | CACCATGGATGGTAAAGGTCAAGATTAC |
| | | R | AATATCACTTCCCTTACCATTAGGCG |
| At4g12020.3 | ID#8 | F | GGAGATAGAACCATGGATTTTTGTATAGCT |
| | | R | CAAGAAAGCTGGGTCTCCATCCCAATCATGTTCA |
| | ID#3 | F | GGAGATAGAACCATGGTTGTAACCAATC |
| | | R | CAAGAAAGCTGGGTCTCCACGGAGGAGCGAAG |
| At4g19500.2 | ID#9 | F | CACCATGACTGGTGTTCTTTCTGTTTTA |
| | | R | AAGTATGCATTCTTTTTCTCGTGTCATATCC |
| At4g12020.1 | ID#4 | F | CACCATGGTTGTAACCAATCCCAATGCC |
| | | R | AAACAGCGGGAAGGATTTTAAAACAATAACCTCT |
| At5G45260.1 | ID#10 | F | CACCATGAAGTGTCTAAATGATAGGTTC |
| | | R | GCAGATGGAGGAGGAAGTGGA |
| At5G47260.1 | ID#11 | F | GGAGATAGAACCATGATAATTCCTCAATTC |
| | | R | CAAGAAAGCTGGGTCGGTATAACATAAAAGGAAATTC |
| Solyc04g008200.2.1 | ID#18 | F | CACCATGTATGCCATCTCTGAATGCTCT |
| | | R | GATCCAGGCTTCTTCTTCCTCATGACCT |
| Solyc05g005130.1.1 | ID#19 | F | CACCATGGCCATGGAGTGTGGCAAAGTG |
| | | R | GTGAAAGCCCTCCATTTCTTCATTTTCTCTTGG |
| Solyc04g007490.2.1 | ID#20 | F | GGAGATAGAACCATGCTGGCATACATAGA |
| | | R | CAAGAAAGCTGGGTCACAGACATCTGTAAAACCAACCA |
| Solyc05g012890.1.1 | ID#21 | F | CACCATGCAGGCAAGCCATTCGTTTTTA |
| | | R | AAAGCCTACTATTTCTTCATTGATCCTTTGTTTACTAGC |
| Solyc10g008220.2.1 | ID#34 | F | CACCATGGCTGATGCTGTTGTCGAATTT |
| | | R | AAAGCCCACCACATTTTCTTCTTCCACCAT |
| Solyc05g013260.1.1 | ID#13 | F | GGAGATAGAACCATGCAATATATTATTGACA |
| | | F | CAAGAAAGCTGGGTCGAAACTCTCCATTTCTTCAT |
| Solyc06g065000.1.1 | ID#32 | F | CACCATGAGAACATTTGTTCTGTTTGGG |
| | | R | GCCAATGATCCAAGCTTCGTCTTCC |
| Solyc05g007170.2.1 | ID#26 | F | CACCATGAAATTGCATGATCTTCTAGTG |
| | | R | GAAAAGCACCATTTCTTCATTTCCTGT |

| | | | |
|---|---|---|---|
| Solyc09g072940.1.1 | ID#27 | F | CACCATGTTTGAATCTACCGAAAGACTT |
| | | R | CGTTGCCAAGATTTTGAAAATAGGCAAG |
| Solyc05g007640.2.1 | ID#15 | F | CACCATGTTTCCAGATAAAGATAATCTC |
| | | R | AAAGCCTACAATATCATCATTAATCGTTT |
| Solyc05g008650.1.1 | ID#23 | F | GGAGATAGAACCATGGAGTGTGCTGACAA |
| | | R | CAAGAAAGCTGGGTCAAAACCAACTACCACTTCATT |
| Solyc05g012910.2.1 | ID#28 | F | GGAGATAGAACCATGATGGAAGATGAATGC |
| | | R | CAAGAAAGCTGGGTCAAAGCCTACGATTTCTTCATT |
| Solyc04g008180.1.1 | ID#16 | F | CACCATGAGTGATCTTATTGATGATTTA |
| | | R | GATCCAGGCTTCTTCTTCCTC |
| Solyc05g012740.1.1 | ID#29 | F | GGAGATAGAACCATGTCGAATTTGTGTAGA |
| | | R | CAAGAAAGCTGGGTCCACGTCCTGAAAGCCCACCAT |
| Solyc05g012740.1.1 | ID#24 | F | CACCATGAAAGAGAAGCCTTCCGAGGAT |
| | | R | ACGACGACCCCAAGAAAACTTGAGCATTT |
| Solyc05g005330.2.1 | ID#17 | F | CACCATGTCTTCTACTTCTTCATATGAA |
| | | R | ATCAAAACCCACCATTTCTTCTCTTG |
| Solyc05g007350.1.1 | ID#30 | F | CACCATGTTTTCCGGCGAATATTTGTCT |
| | | R | AAAACCAACAACTTCTTCATCGGTTATCCTTGGA |
| Solyc10g051170.1.1 | ID#25 | F | GGAGATAGAACCATGGGGACTCTTGTACAC |
| | | R | CAAGAAAGCTGGGTCATCAAAACCCACCATTTCTT |
| Solyc01g102880.1.1 | ID#31 | F | GGAGATAGAACCATGAAGATCCAGGGCA |
| | | R | CAAGAAAGCTGGGTCGTTATTATCCCACCAAAACT |
| Medtr3g030980.1 | ID#36 | F | GGAGATAGAACCATGGATGATTGGAGAAGTCTGG |
| | | R | CAAGAAAGCTGGGTCTATTGTAACCTGGTCATTTAT |
| Medtr3g015260.2 | ID#37 | F | GGAGATAGAACCATGTCCATGGCAGAGTTGGTT |
| | | R | CAAGAAAGCTGGGTCGTCACCGTTTCTACCATAT |
| Medtr3g032760.1 | ID#38 | F | GGAGATAGAACCATGAGGAGTCTGGATTCCATTT |
| | | R | CAAGAAAGCTGGGTCTATTGTGACTTGGTCGTTTA |
| Medtr3g018930.2 | ID#40 | F | GGAGATAGAACCATGTCCATGAGAGAGTTGGTTG |
| | | R | CAAGAAAGCTGGGTCATCTCTACCATATATGCTCG |
| Medtr6g087200.1 | ID#41 | F | GGAGATAGAACCATGGAACGTTGCAGTAGCTTG |
| | | R | CAAGAAAGCTGGGTCAACAATCCTTGATCTAATTTG |
| Medtr6g015490.1 | ID#42 | F | GGAGATAGAACCATGGACAAGCTAAAAGCAAAGG |
| | | R | CAAGAAAGCTGGGTCCACTTCATAAAGTCTATTTTC |
| Medtr6g015665.1 | ID#43 | F | GGAGATAGAACCATGGGAAAGAAAAACGACGAAC |
| | | R | CAAGAAAGCTGGGTCCACTTCATGAAGTTTATTTTC |
| MDP0000286727 | ID#52 | F | GGAGATAGAACCATGGGAATTTTTCTCTCTGGAAAT |
| | | R | CAAGAAAGCTGGGTCGCGTGACCTTCTGTAAAAG |
| MDP0000457940 | ID#88 | F | GGAGATAGAACCATGGAAGCTTGCCATTCACTTG |
| | | R | CAAGAAAGCTGGGTCATGAGCTCCTGGTGCATC |
| Bradi4g24914 | ID#45 | F | GGAGATAGAACCATGGCAACACATGCAACTACAC |
| | | R | CAAGAAAGCTGGGTCCAGATCACAGTTTGCATTAA |

| | | | |
|---|---|---|---|
| Bradi3g34961.1.p | ID#55 | F | GGAGATAGAACCATGGAGGATGCCAATATAGAGG |
| | | R | CAAGAAAGCTGGGTCTTCCTCAAGCACTTCGAG |
| Bradi3g58937.5.p | ID#64 | F | GGAGATAGAACCATGGGAGAAAGCTTGTGGGAAA |
| | | R | CAAGAAAGCTGGGTCGTGACAAACATCCAATGCTT |
| LOC_Os02g19890.3 | ID#67 | F | GGAGATAGAACCATGCGACAGGATGATCTTCGTA |
| | | R | CAAGAAAGCTGGGTCCATAGGGTGAAAATAGACCC |
| LOC_Os11g11920.1 | ID#68 | F | GGAGATAGAACCATGGGGAACAGTATGGTGATA |
| | | R | CAAGAAAGCTGGGTCTTTGCACCATCCAAAGCAG |
| LOC_Os07g17230.1 | ID#70 | F | GGAGATAGAACCATGGAACCTGGAAGCAGCAGT |
| | | R | CAAGAAAGCTGGGTCAGAGTAGTTCAGATCATCAAA |
| LOC_Os05g15040.1 | ID#71 | F | GGAGATAGAACCATGGTAAATGAGGCCTGGAAGG |
| | | R | CAAGAAAGCTGGGTCTCCGCTGTCACCAAAACG |
| LOC_Os12g18360.1 | ID#72 | F | GGAGATAGAACCATGGCGAATGAGGGGTTCGG |
| | | R | CAAGAAAGCTGGGTCACGGAGGAAGTAAGGAGAA |
| LOC_Os08g30634.1 | ID#73 | F | GGAGATAGAACCATGGCTATGCCCAACCTCCAA |
| | | R | CAAGAAAGCTGGGTCCCTGGTATCGGTTTCTTTC |
| LOC_Os09g20020.1 | ID#74 | F | GGAGATAGAACCATGGATTCTGGTAATATTGGAGG |
| | | R | CAAGAAAGCTGGGTCAGAAAACTCTGGGCATGAAG |
| LOC_Os11g11810.1 | ID#75 | F | GGAGATAGAACCATGGTTATTGTGAAGATGCCCAC |
| | | R | CAAGAAAGCTGGGTCAAATCGTCCACCTCTCTTG |
| LOC_Os11g46210.1 | ID#76 | F | GGAGATAGAACCATGCAAGGGCTAACCGAAGAG |
| | | R | CAAGAAAGCTGGGTCTACAAGCATTTGACATATCTC |
| Glyma.12G236500.1.p | ID#83 | F | GGAGATAGAACCATGGGTCGACCTAGAGGTGA |
| | | R | CAAGAAAGCTGGGTCTAACTCAAATTGTTCACCAAC |
| Glyma.05G165800.4.p | ID#85 | F | GGAGATAGAACCATGTATCTTCCTGAATTGGTCAA |
| | | R | CAAGAAAGCTGGGTCTAACTTGCAGGATTGACTTGC |
| Glyma.05G165800.4.p | WRKY-ID#85 | F | GGGGACAAGTTTGTACAAAAAAGCAGGCTTAATGGCTGAGCCAAAGATATTTC |
| | | R | GGGGACCACTTTGTACAAGAAAGCTGGGTAGTCTAACTTGCAGGATTGACTT |
| Glyma.05G165800.4.p | GmNLR-ID#85 | F | GGGGACAAGTTTGTACAAAAAAGCAGGCTTAATGGCTTCCTCGGCCTTGA |
| | | R | GGGGACCACTTTGTACAAGAAAGCTGGGTCTAACTTGCAGGATTGACTTGCCTCA |
| At4g12020.3 | ID#3A | F | GGAGATAGAACCATGGTTGTAACCAATC |
| | ID#3A | R | CAAGAAAGCTGGGTCATCCTTCGGGTTAAGAATAA |
| At4g12020.3 | ID#3B | F | GGAGATAGAACCATGTATGATGGTTATGGAACTCC |
| | ID#3B | R | CAAGAAAGCTGGGTCTCCACGGAGGAGCGAAG |
